# Synapse type-specific molecular nanoconfigurations of the presynaptic active zone in the hippocampus identified by systematic nanoscopy

**DOI:** 10.1101/2022.03.11.483942

**Authors:** Hirokazu Sakamoto, Naoya Kimpara, Shigeyuki Namiki, Shun Hamada, Toshihisa Ohtsuka, Kenzo Hirose

**Affiliations:** Department of Pharmacology, Graduate School of Medicine, The University of Tokyo, Tokyo 113-0033, Japan; Department of Biochemistry, Faculty of Medicine, University of Yamanashi, Yamanashi, Japan; International Research Center for Neurointelligence (WPI-IRCN), The University of Tokyo, Tokyo, 113-0033, Japan

**Keywords:** synapse, neurotransmitter release, active zone, synaptic diversity, super- resolution imaging

## Abstract

Neurotransmitters are released by exocytosis from synaptic vesicles at the active zone in the presynaptic terminal. The scaffold of the active zone consists of only a few evolutionarily conserved proteins, including RIM, CAST/ELKS, and RIM-BP, and tethers Munc13 and Ca^2+^ channels. The molecular principles that enable these proteins to mediate synaptic diversity have remained unclear. Here, we identified synapse type-specific molecular nanoconfigurations in the active zone by systematic quantification of active zone proteins using nanoscopy at two types of excitatory synapses and two types of inhibitory synapses in the rat hippocampal CA3 region. Quantitative analysis revealed that Munc13 content was particularly varied among the various synapse types and that the physical proximity of Ca^2+^ channels to the active zone scaffolds correlated with the efficacy of neurotransmitter release. We propose that the active zone is a flexible supramolecular assembly that can tune its composition and spatial configuration to adjust neurotransmitter release.

## Introduction

Chemical synaptic transmission is fundamental to information processing in the brain. Typically, neurotransmitters are released by exocytosis from synaptic vesicles at a specialized membrane domain of the presynaptic terminal known as the active zone (*1, 2*). The active zone contains proteinaceous materials that dock and tether synaptic vesicles to the plasma membrane and is precisely opposed to the postsynaptic domain where neurotransmitter receptors accumulate (*3*). The core of the mammalian active zone consists of a few evolutionarily conserved proteins, including RIM, CAST/ELKS, RIM-BP, and Munc13 (*4, 5*). Genetic deletion of Munc13 leads to total arrest of synaptic vesicle exocytosis (*6–8*), and to loss of docked synaptic vesicles (*8–10*). Munc13 makes synaptic vesicles release-ready at spatially restricted sites within the active zone by specifying the site of docking/priming by recruiting a t-SNARE protein, syntaxin-1 (*11*). Interestingly, combined, but not individual, genetic deletion of RIM and CAST/ELKS leads to loss of Munc13 and other active zone proteins (*12*). Also, combined deletion of RIM and RIM-BP leads to a severe reduction of Munc13 (*13*). At synapses in these combined deletion mutants, Munc13 loss is accompanied by a reduction in Ca^2+^ channel tethering at the active zone with a concomitant drastic reduction in neurotransmitter release. Therefore, RIM, CAST/ELKS, and RIM-BP are vital but redundant constituents of the active zone scaffold that tethers the release machinery, Munc13 and Ca^2+^ channels, at the presynaptic membrane.

Diversity of synaptic properties is fundamental to brain function, but the molecular basis of this diversity remains elusive. Although synapses formed by distinct types of neurons in the brain share similar kinds of synaptic proteins, they display very diverse characteristics of neurotransmitter release, such as release probability and short-term plasticity (*14–16*). The hippocampal CA3 region is an attractive brain region to explore synaptic diversity because of its simple synaptic organization and its intelligible roles in learning and memory (*17*). Furthermore, electrophysiological properties of principal glutamatergic (*18–20*) and GABAergic (*21–23*) synapses in this region have been well characterized. Freeze-fracture immunogold labeling has been applied to analyze the nanoscale distribution of active zone proteins and Ca^2+^ channels in the CA3 region (*24, 25*) and recently to analyze synapse type-specific molecular distributions correlating to synaptic properties (*26–29*). Super-resolution immunohistochemical fluorescence imaging of synaptic proteins at various types of physiologically characterized synapses is a promising alternative approach to investigate the molecular basis of synaptic diversity (*30–32*). In this study, we systematically determined the composition and the nanoscale spatial distribution of active zone molecules at two major glutamatergic synapses, associative/commissural fiber (AC) and mossy fiber (MF) synapses, as well as at two types of perisomatic GABAergic basket cell synapses in the rat hippocampal CA3 region using multicolor super-resolution fluorescence imaging techniques. These methods are suitable for analyzing sub-synaptic structures (*33, 34*) and to determine the molecular correlates of functional properties of synapses (*11*). Our findings reveal synapse type-specific molecular nanoconfigurations of the active zone that account for the diversity of neurotransmitter release efficacy among synapses.

## Results

### Immunohistochemistry of active zone proteins in hippocampal synapses

We first visualized seven proteins *in situ*, five active zone proteins (Munc13, RIM1, RIM2, CAST, and RIM-BP2) and two Ca^2+^ channel subunits (Cav2.1 and Cav2.2) in the rat hippocampus by immunohistochemistry (Fig. 1). At low magnification, the staining pattern of an antibody recognizing all isoforms of Munc13 was consistent with the laminar synaptic organization of the hippocampus and essentially identical to that of an antibody specific to Munc13-1 (Fig. 1 A and B). Antibodies against the other six proteins also showed laminar staining patterns in the hippocampus with unique laminar preferences (Fig. S1 A–G). To compare the expression profile of these proteins among strata, the signal intensity of each protein was normalized against Munc13 signal intensity (Fig. S1 A–F bottom panels, and H). In the CA3 region, the relative intensities of CAST, RIM-BP2, and Cav2.1 were lower in the stratum lucidum, where MF axons from dentate gyrus granule cells terminate, than in the other strata. In the inner molecular layer of the dentate gyrus, the relative intensities of RIM1, RIM2, and CAST were particularly high. The relative intensity of Cav2.2 was markedly low in the strata where perforant path axons from the entorhinal cortex terminate (i.e., lacunosum-moleculare of both CA3 and CA1, as well as middle to outer molecular layers of the dentate gyrus). At high magnification, Munc13 staining showed small punctate patterns and totally matched that of Munc13-1 (Fig. 1C). The staining of the other six proteins also showed punctate patterns, and closely resembled that of Munc13 (Fig. 1 D–I). Thus, these punctate signals should correspond to the presynaptic active zones.

**Figure 1.**
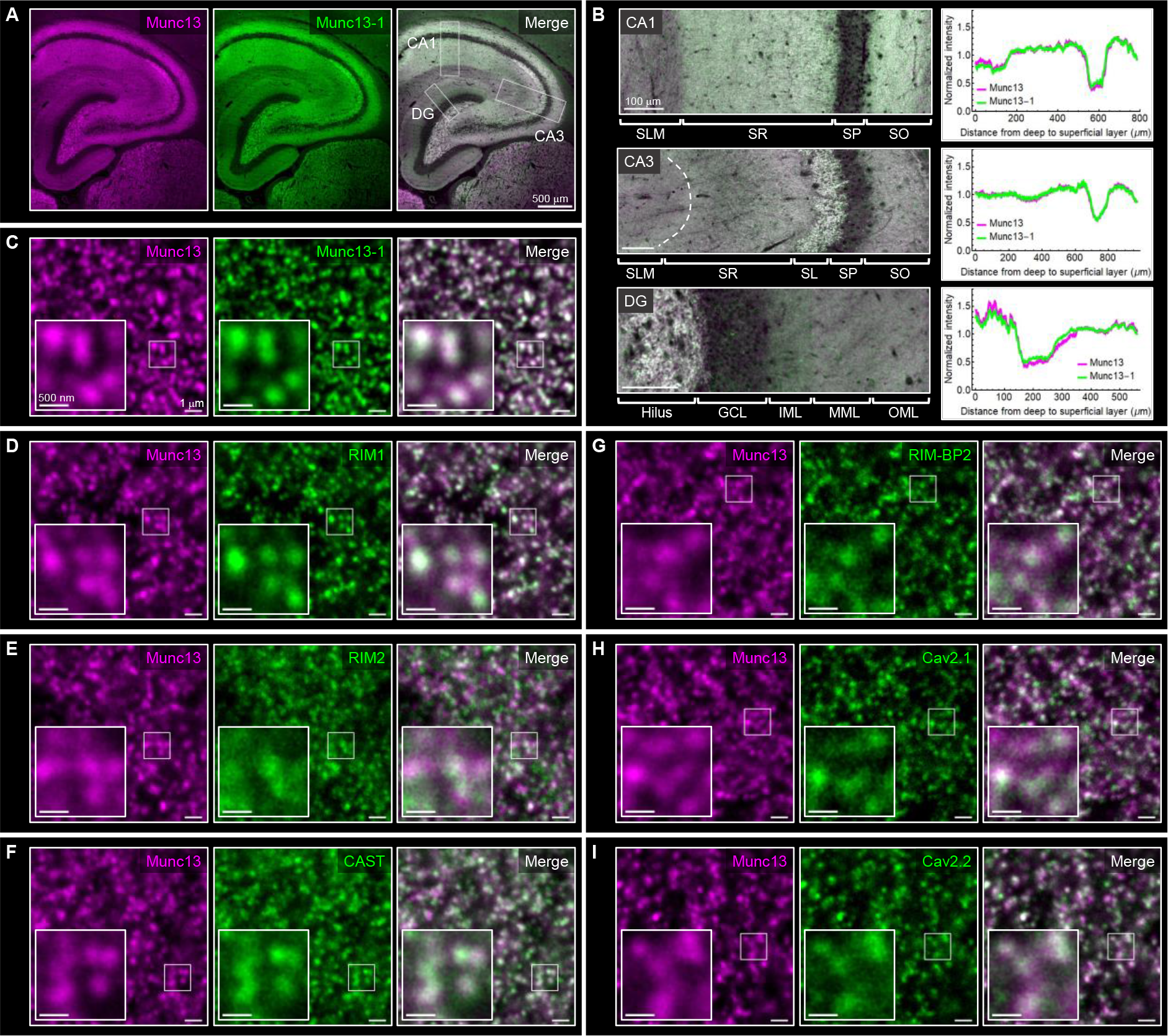
Immunohistochemical analysis of active zone proteins in the hippocampus. (***A***) Immunofluorescence images of Munc13 (magenta) and Munc13-1 (green) in the rat hippocampus at low magnification. Scale bar represents 500 µm. (**B**) Enlarged views of white boxed regions in (A) and corresponding intensity profiles of Munc13 and Munc13-1 immunofluorescence in CA1, CA3, and dentate gyrus (DG). Scale bars represent 100 µm. (**C**–**I**) High magnification images of Munc13 (magenta) and active zone proteins (green), Munc13-1 (C), RIM1 (D), RIM2 (E), CAST (F), RIM- BP2 (G), Cav2.1 (H), and Cav2.2 (I), in the stratum radiatum of the hippocampal CA3 region. Scale bars: 1 µm for the original panels and 500 nm for the enlarged panels, respectively.

To characterize active zones of glutamatergic and GABAergic synapses in the hippocampal CA3 region in more detail, we simultaneously imaged Munc13 with synapse-type specific markers at synapse resolution (Fig. 2 A–D). In the stratum radiatum, 82 ± 3% of Munc13 puncta were positive for PSD95, a postsynaptic marker for glutamatergic synapses (Fig. 2 B and E), and almost all of them were positive for VGLUT1 (Fig. S2A). Furthermore, most Munc13 puncta in this stratum were positive for GluA2, which is expressed at high levels in CA3 pyramidal cells (*35*), and a tiny fraction of Munc13 puncta were strongly positive for GluA4 or mGluR1α, which are expressed at high levels in specific subsets of interneurons (Fig. S2 B and C). In the stratum lucidum, 72 ± 4% of Munc13 puncta were positive for PSD95 (Fig. 2 C and E) and were highly clustered in large VGLUT1 positive terminals (Fig. S2A), which is a hallmark of MF synapses onto CA3 pyramidal cells. In contrast to other strata, a large fraction of Munc13 puncta in the stratum pyramidale was positive for gephyrin, a postsynaptic marker for GABAergic synapses (Fig. 2 D and E), indicating that these puncta correspond to GABAergic inputs of perisomatic basket cells. Perisomatic basket cells consist of two major types of interneurons, parvalbumin (PV)-containing fast-spiking interneurons and cholecystokinin (CCK)-containing regular-spiking interneurons (*36*). The former specifically express synaptotagmin 2 (Syt2) for fast- synchronous release (*37*), and a large fraction of the latter expresses VGLUT3 (*38*). Indeed, 60 ± 3% and 34 ± 4% of GABAergic terminals identified by VGAT were positive for Syt2 and VGLUT3, respectively, in the stratum pyramidale of the CA3 (Fig. S2 D and E). These two presynaptic proteins were distributed in a mutually exclusive manner (Fig. 2F); therefore, we concluded that Syt2- and VGLUT3-positive GABAergic active zones were inputs from PV- and CCK-containing basket cells, respectively.

**Figure 2.**
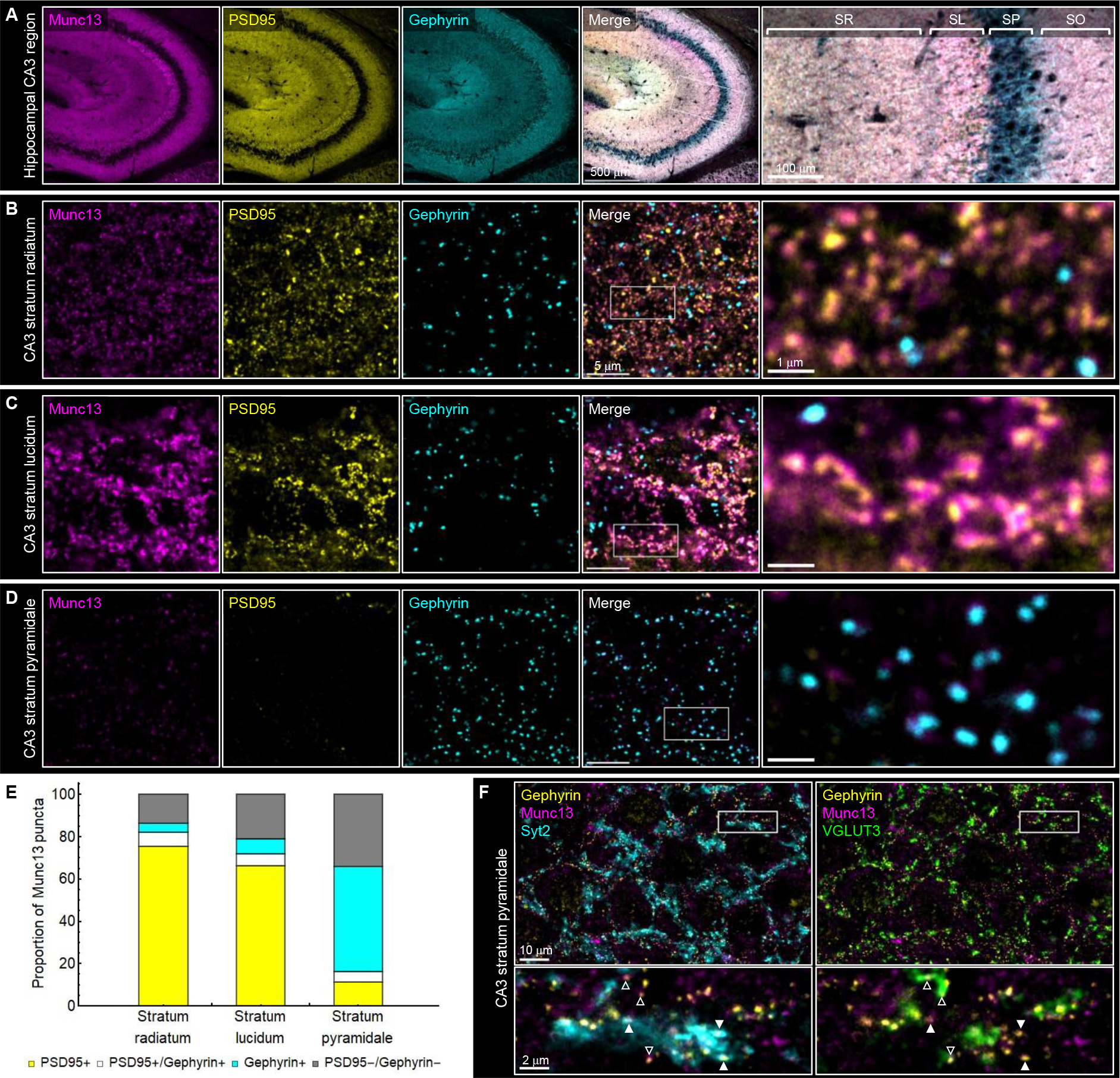
Characterization of glutamatergic and ABAergic active zones in the hippocampal CA3 region. (**A**) Immunofluorescence images of Munc13 (magenta), PSD95 (yellow), and gephyrin (cyan) in the hippocampal CA3 region at low magnification. The rightmost panel represents an enlarged view of the white boxed region in the merged image. Scale bars: 500 µm for the original images and 100 µm for the enlarged image. (**B**–**D**) High magnification images of Munc13 (magenta), PSD95 (yellow), and gephyrin (cyan) in the stratum radiatum (B), the stratum lucidum (C), and the stratum pyramidale (D) of the hippocampal CA3 region. The rightmost panels represent an enlarged view of the white boxed region in the merged image. Scale bars: 5 µm for the original images and 1 µm for the enlarged images. (**E**) Summary of the ratio of Munc13 puncta that are positive for PSD95 or gephyrin in the stratum radiatum (*n* = 4,359 puncta from 3 animals), the stratum lucidum (*n* = 3,004 puncta from 3 animals), and the stratum pyramidale (*n* = 1,134 puncta from 3 animals). (**F**) Four color immunofluorescence images of gephyrin (yellow), Syt2 (cyan), G UT3 (green), and Munc13 (magenta) in the stratum pyramidale of the hippocampal CA3 region. Bottom panels represent enlarged views of the white boxed region in the top images. Syt2 and G UT3 distribute in a mutually exclusive manner (closed and open triangles). Scale bars: 10 µm for the original images and 2 µm for the enlarged images.

### Super-resolution imaging of Munc13 at synapses in the hippocampal CA3 region

We have previously shown that Munc13-1 forms nanosized supramolecular assemblies, which correspond to synaptic vesicle release sites at glutamatergic synapses in cultured hippocampal neurons using three-dimensional stochastic optical reconstruction microscopy (STORM) (*11*). To ask whether similar supramolecular assemblies are observed in native synapses in the hippocampus, we performed STORM imaging of Munc13 with postsynaptic markers (PSD95 or gephyrin) on immunostained sections. Two-color STORM clearly resolved nanosized (∼100 nm) assemblies of Munc13 aligned with PSD95 scaffolds in the stratum radiatum (Fig. 3A) and stratum lucidum (Fig. 3B). Also, in the stratum pyramidale, we found nanosized assemblies of Munc13 located close to gephyrin scaffolds (Fig. 3C). Thus, we concluded that Munc13 consistently forms nanosized assemblies at both glutamatergic and GABAergic synapses in the hippocampus.

**Figure 3.**
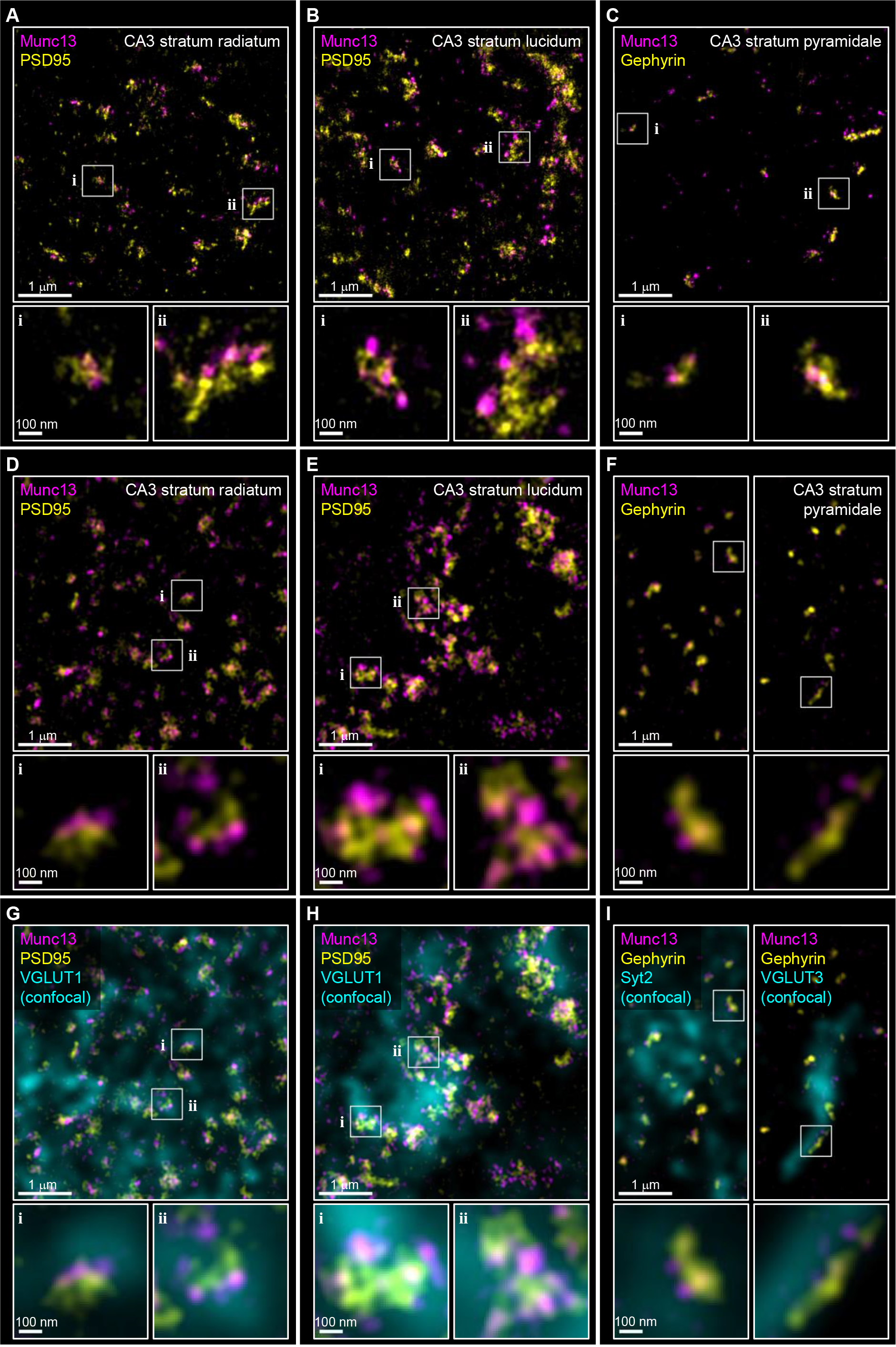
Nanoscopic imaging of Munc13 at synapses in the hippocampal CA3 region. (**A**–**C**) Two-color STORM images of Munc13 (magenta) and postsynaptic scaffolds (PSD95 or gephyrin; yellow) in the stratum radiatum (A), stratum lucidum (B), and stratum pyramidale (C). Bottom panels show enlarged views of white boxed regions in the top panels. (**D**–**F**) Two-color STED images of Munc13 (magenta) and postsynaptic scaffolds (PSD95 or gephyrin; yellow) in the stratum radiatum (D), stratum lucidum (E), and stratum pyramidale (F). Bottom panels show enlarged views of white boxed regions in the top panels. (**G** and **H**) STED images in (D) and (E) overlaid with a confocal channel for an additional excitatory presynaptic marker, VGLUT1. (**I**) STED images in (F) overlaid with a confocal channel for additional presynaptic markers: Syt2 (left) and VGLUT3 (right). The scale bars for top and bottom images represent 1 µm and 100 nm, respectively.

Consistent with the results obtained by STORM, we confirmed that nanosized assemblies of Munc13 aligned with postsynaptic scaffolds (PSD95 or gephyrin) were resolved in brain sections by two-color stimulated emission depletion (STED) microscopy when the depletion laser power was sufficiently elevated (Fig. 3 D–F; see Materials and Methods). STORM provides superior spatial resolution compared with STED microscopy both in lateral and axial directions but generates a large volume of raw image data and is time-consuming. Our STED analysis sufficiently resolved sub- synaptic Munc13 nanoassemblies; therefore, we decided to use STED microscopy in subsequent nanoscopy experiments to perform a systematic survey. Furthermore, the use of STED microscopy enabled the synapse type to be determined by simultaneously imaging presynaptic markers (VGLUT1, VGLUT3, or Syt2) using a confocal channel (Fig. 3 G–I).

### Molecular composition of the active zone at multiple types of synapses in the hippocampal CA3 region

To understand the diversity of active zone molecular architectures among the four types of synapses in the hippocampal CA3 region, we performed a systematic nanoscopic analysis of the seven proteins (Munc13, RIM1, RIM2, CAST, RIM-BP2, Cav2.1, and Cav2.2). All pair combinations of the seven proteins were imaged using two-color STED channels (21 combinations) with imaging of PSD95 by a confocal channel for two types of glutamatergic synapses (Figs. 4 and S3) or with imaging of gephyrin and presynaptic markers (Syt2 or VGLUT3) by two confocal channels for two types of GABAergic synapses (Figs. 5 and S4). STED microscopy revealed that, similar to Munc13 (see Fig. 3), the other six proteins also showed nanosized (∼100 nm) sub-active zone hotspots at all types of synapses examined.

**Figure 4.**
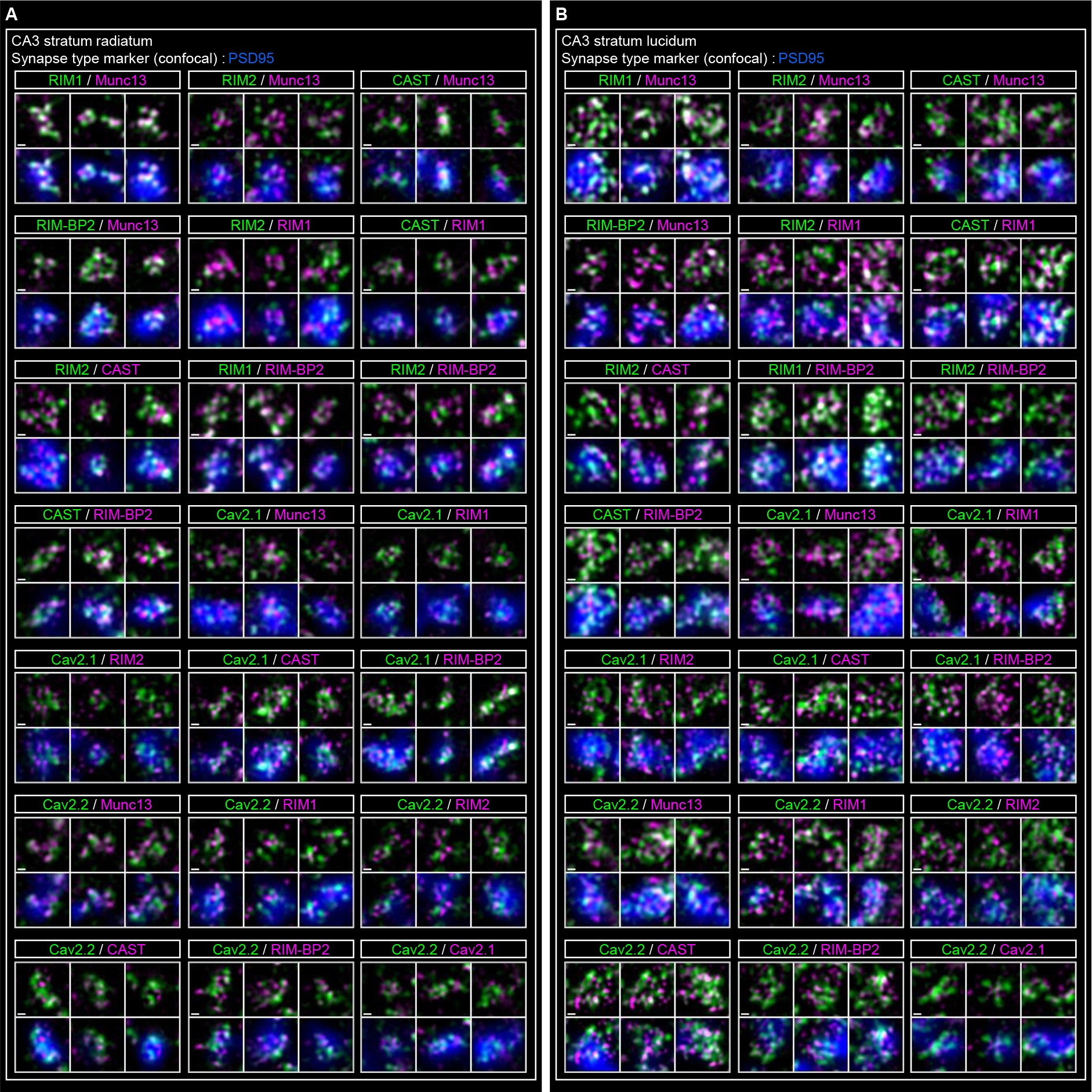
Systematic nanoscopic imaging of active zone proteins at glutamatergic synapses in the hippocampal CA3 region. (**A** and **B**) Two-color STED images of active zone proteins in AC synapses in the stratum radiatum (A) and MF synapses in the stratum lucidum (B). For each synapse type, all 21 combinations of protein pairs for the seven active zone proteins were subjected to two-color STED imaging. The resultant STED images (magenta and green) are shown with (bottom) and without (top) overlay of the confocal channel for PSD95 (blue), which identifies excitatory synapses in both regions. Scale bars, 100 nm.

**Figure 5.**
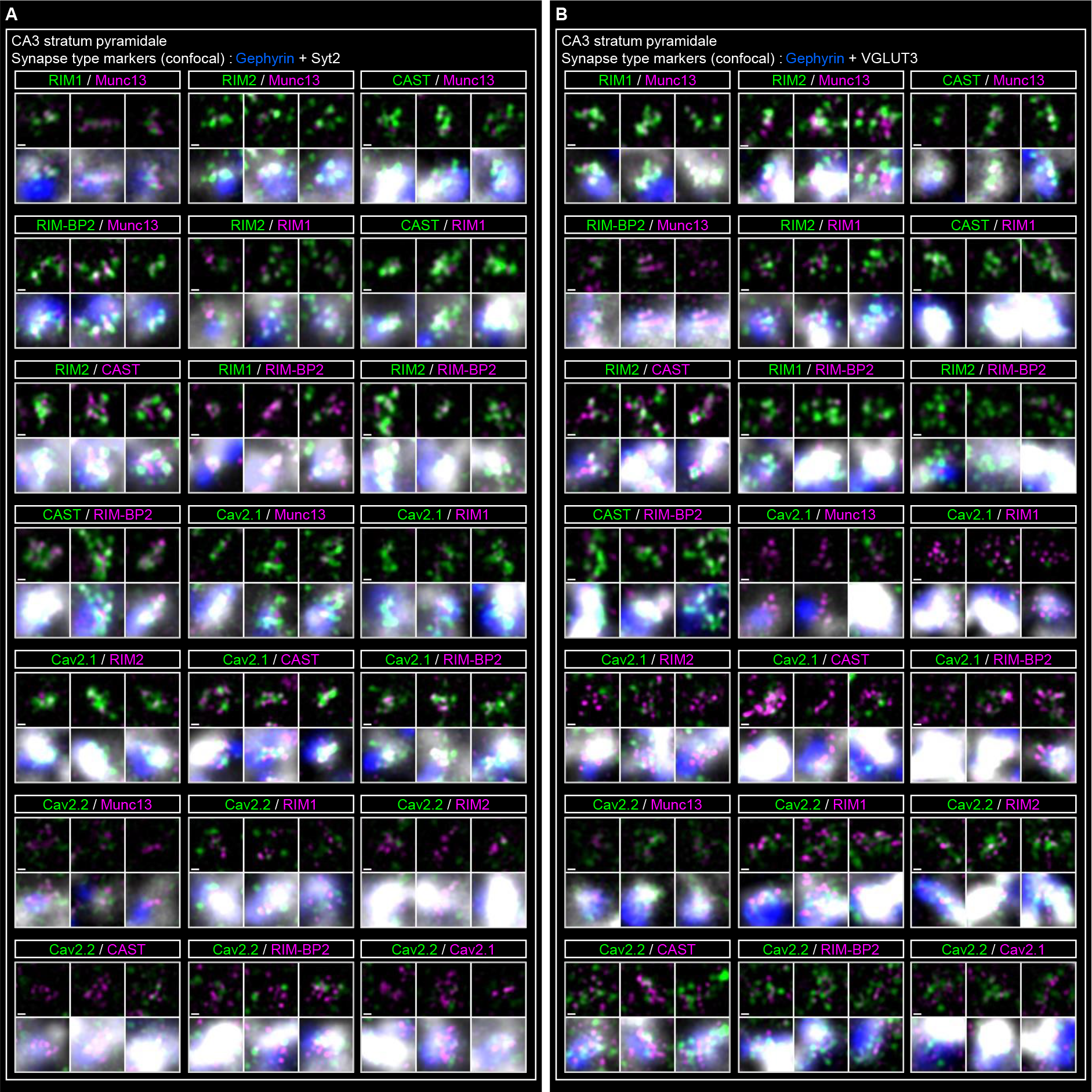
Systematic nanoscopic imaging of active zone proteins at perisomatic ABAergic synapses in the hippocampal CA3 region. (**A** and **B**) Two-color STED images of pairs of active zone proteins in P synapses (A) and CCK synapses (B) in the stratum pyramidale. For each synapse type, all 21 combinations of protein pairs for the seven active zone proteins were subjected to two-color STED imaging. The resultant STED images (magenta and green) are shown with (bottom) and without (top) overlay of the confocal channels for gephyrin (blue) and a synapse type-specific marker (white). Syt2 and G UT3 are synapse type-specific markers to identify P and CCK synapses, respectively. Scale bars, 100 nm.

We constructed a nanoscopic image processing pipeline to quantitatively analyze the composition and distribution of proteins in the active zone of a large population of synapses (Fig. 6). First, image masks for the active zones were generated from merged images of two-color STED channels by unsharp masking, which defined the area of individual active zones. Second, image masks for sub-active zone hotspots for individual proteins were separately generated from respective STED channels by image deconvolution. Active zones were then classified into glutamatergic or GABAergic synapses by co-localization with synapse type-specific markers (PSD95 or gephyrin, respectively) imaged by a confocal channel. Identified glutamatergic active zones in the stratum radiatum and the stratum lucidum were further categorized as AC- and MF synapses, respectively (see Fig. 2). An additional confocal channel of a presynaptic marker (Syt2 or VGLUT3) was used to categorize GABAergic active zones in the stratum pyramidale as PV (Gephyrin+/Syt2+) and CCK (Gephyrin+/VGLUT3+) synapses, respectively. The obtained masks for active zones were used to calculate the density of respective proteins and the spatial correlation between the pair of proteins, while those for sub-active zone hotspots were used to quantify the number and size of hotspots, the local density of the protein, and the degree of co-localization (see below).

**Figure 6.**
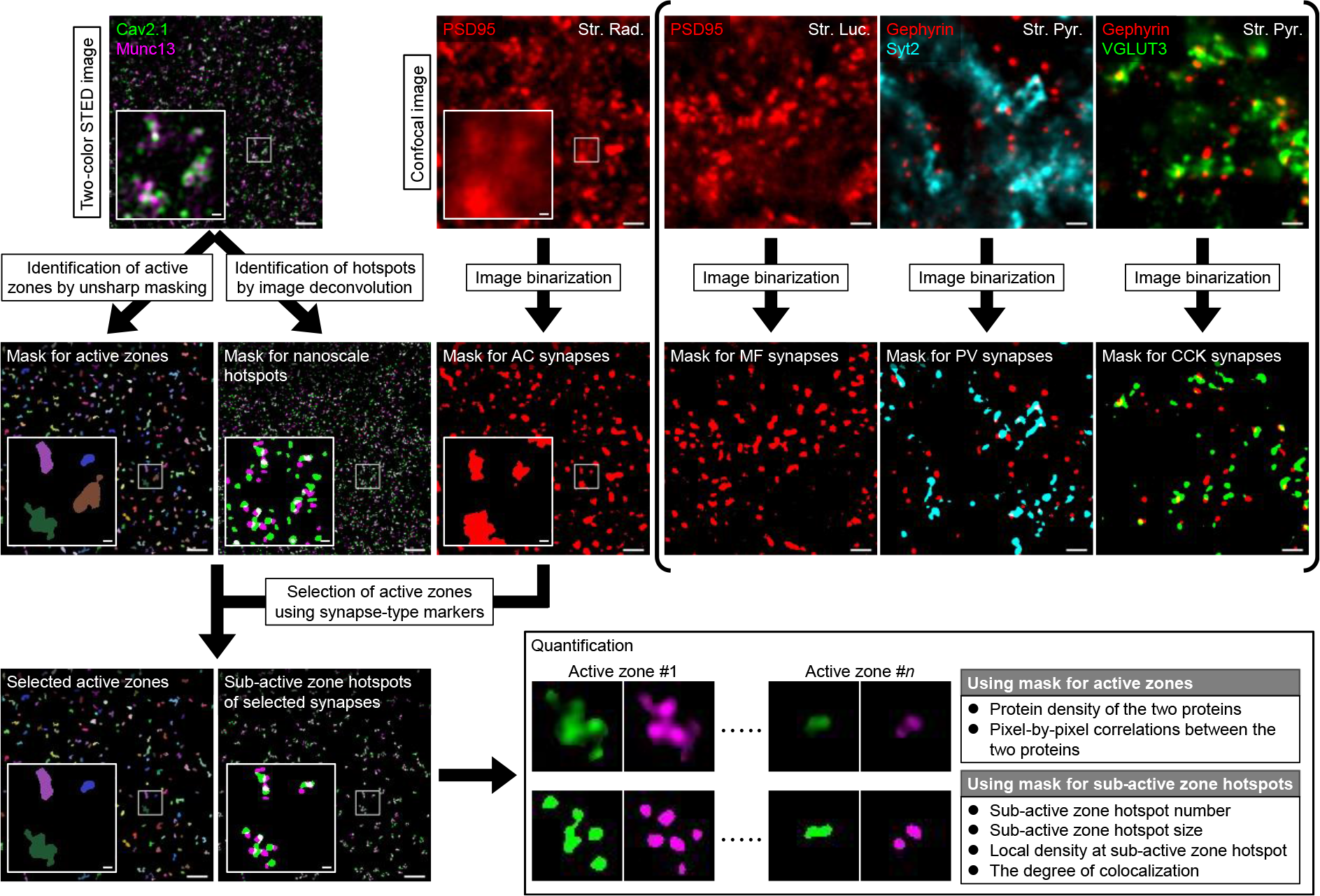
Workflow of systematic nanoscopic analysis of active zone proteins. A systematic nanoscopic image processing pipeline for quantitative analysis of the composition and distribution of proteins in the active zone is shown. Image masks for active zones and sub-active zone hotspots are generated from two-color STED images, by unsharp masking and image deconvolution, respectively. Each active zone and corresponding sub-active zone hotspots are then classified into defined types of synapses by a binarized mask of confocal channels detecting synapse type-specific markers: PSD95 for AC and MF synapses, gephyrin and Syt2 for PV synapses, and gephyrin and VGLUT3 for CCK synapses. Obtained masks for active zones covering the total active zone area are used to calculate the density of respective proteins and spatial correlation between a pair of proteins. Masks for sub-active zone hotspots are used to quantify the number and size of hotspots, the local density of the protein, and the degree of co-localization of a pair of proteins. Scale bars: 1 µm for the original images and 100 nm for the enlarged views.

As a result, we identified a total of 61,456 active zones and quantified the normalized density at the active zone (Fig. 7A), and the number (Fig. 7B), size (Fig. 7C) and local density (Fig. 7D) of the sub-active zone hotspots of the seven active zone proteins for hundreds to thousands of active zones of the four types of synapses. We found a large difference in the density of Munc13 among the four types of synapses (Fig. 7A). Perisomatic GABAergic synapses possessed low levels of Munc13 compared with glutamatergic synapses, albeit both glutamatergic and GABAergic synapses require Munc13 for synaptic vesicle exocytosis^5^. Notably, the density of Munc13 at PV synapses was only 40% of that at MF synapses. A different set of experiments using an antibody specific to Munc13-1 showed an almost identical trend (Fig. S5A). The number and size of sub-active zone Munc13 hotspots were similar among the four types of synapses (Fig. 7 B and C); therefore, the difference in the overall density of Munc13 may come from the difference in the local density of Munc13 at sub-active zone hotspots (Fig. 7D). Similarly, the density of RIM1 was especially low at PV synapses, while that of RIM-BP2 was low at CCK synapses. Consistent with previous studies (*37, 39*), the density of Cav2.1 was higher at PV synapses than at CCK synapses, and *vice versa* for Cav2.2. The difference in the overall density of Cav2.1 was accounted for by the local density at the sub-active zone hotspots as well as by the size of sub-active zone hotspots.

**Figure 7.**
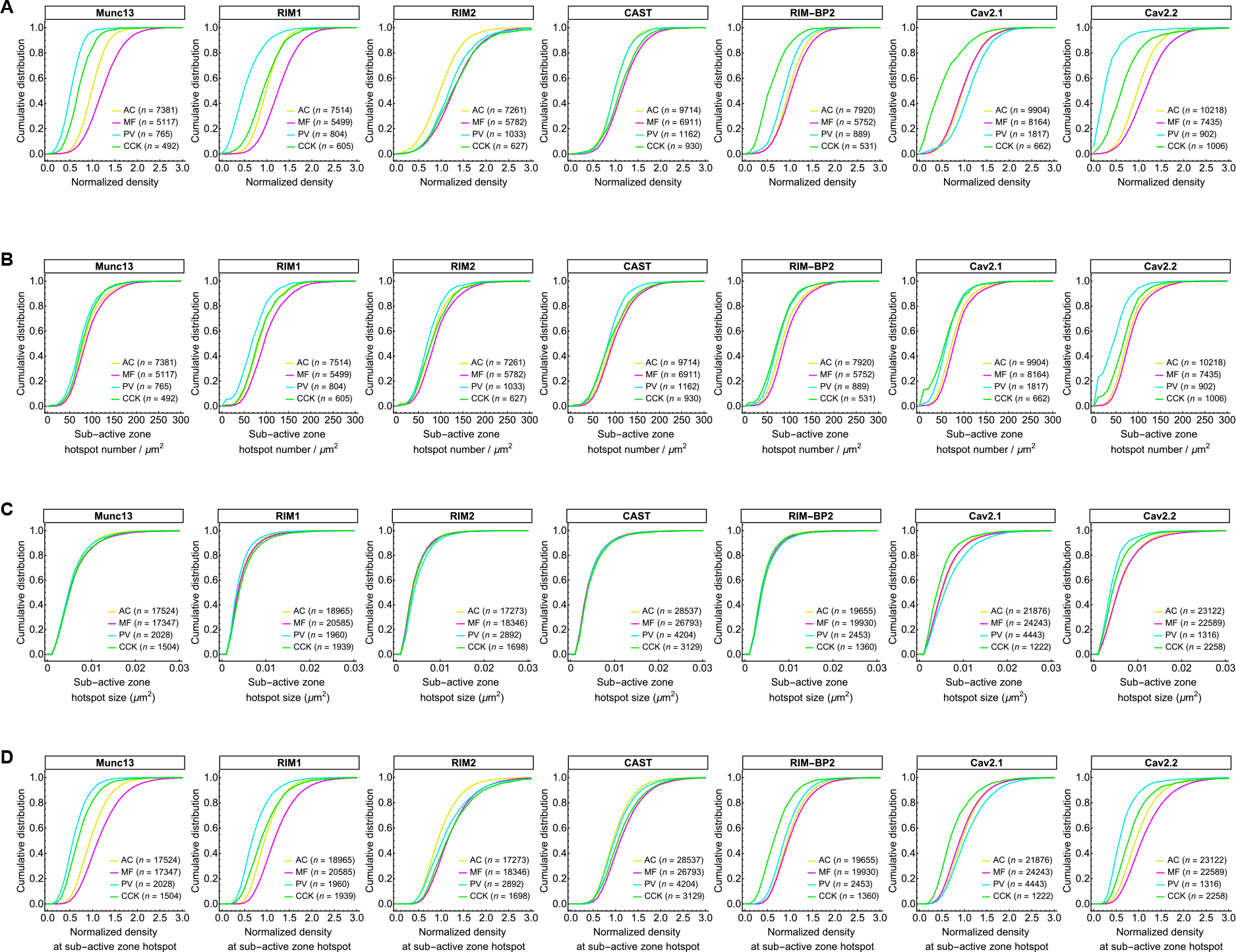
Systematic quantification of active zone proteins at four types of synapses in the hippocampal CA3 region. (**A**) Cumulative plots of the density of active zone proteins for the four types of synapses. The data were normalized against the mean values of AC synapse density for each protein. (**B**) Cumulative plots of the number of sub-active zone hotspots per area for the four types of synapses. (**C**) Cumulative plots of the size of sub-active zone hotspots for the four types of synapses. (**D**) Cumulative plots of the local density of active zone proteins within sub-active zone hotspots for the four types of synapses. The data were normalized against the mean values of AC synapse density for each protein.

Considering that Munc13 is a central component of the synaptic vesicle release site (*11, 40, 41*), it is important to elucidate the quantitative relationship between Munc13 and other active zone proteins. For this purpose, we compared the density of Munc13 and the other proteins in the same active zones (Fig. S5 B–G). The density of Munc13 was lower in the two types of GABAergic synapses than in glutamatergic synapses. However, the densities of the four active zone scaffold proteins were not always lower in these GABAergic synapses: the relative content of RIM2, CAST, and RIM-BP2 was higher in PV synapses, and that of RIM1, RIM2 and CAST was higher in CCK synapses. The density ratio of Cav2.1 to Munc13 differed substantially among synapse types. Notably, the relative content of Cav2.1 to Munc13 was high at PV synapses. In contrast, Cav2.2 scaled linearly with Munc13 across synapse types.

### Distinct clustering of active zone proteins at multiple types of synapses in the hippocampal CA3 region

To further understand the molecular basis of synaptic diversity, we focused on nanoscale inter-molecular spatial relationships between the active zone proteins. In nanometer scale STED images, the seven active zone proteins showed unique spatial distributions (Figs. 4 and 5). Some protein pairs (e.g., RIM1-Munc13) showed a high degree of co-localization, while other protein pairs (e.g., RIM2-Cav2.2) showed almost complete segregation. To quantitatively evaluate spatial relationships, we systematically analyzed the spatial correlations in two ways. First, Pearson’s correlation coefficients (pixel-by-pixel correlations) between two-color STED images of active zone proteins were calculated at individual active zones (Fig. S6). Second, binary masks for sub-active zone hotspots (see Fig. 6) of one protein were used to calculate the ratio of signal intensity between the masked region and the remaining active zone region of the other active zone protein (Fig. S7; see Materials and Methods). In both analyses, a positive value indicates nanoscale clustering of the two proteins, and conversely a negative value indicates segregation of the two proteins at the sub-active zone level. To validate this, we applied these analyses to two-color STED images of Munc13 and Munc13-1. These gave high positive values (∼0.5) at all types of synapses (Figs. S6 and S7), as expected with their overlapping recognition. The two types of independent estimates of spatial correlation were closely matched across protein pairs (Fig. S8); therefore, we treated their mean value as an index of spatial correlation (ISC).

The ISC values differed widely between pairs of the seven proteins, and even between the same protein pairs in different synapse types (ranging from -0.24 to 0.31; Fig. 8 A and B). Notably, a wide variety in ISC values was evident for the four active zone scaffold proteins for each synapse type, indicating that scaffold proteins do not share a common spatial distribution within the active zone. Strong clustering was observed in RIM1-RIM-BP2 pairs in AC-, MF-, and CCK synapses, whereas RIM-BP2 clustered more strongly with CAST and RIM2 than with RIM1 in PV synapses.

**Figure 8.**
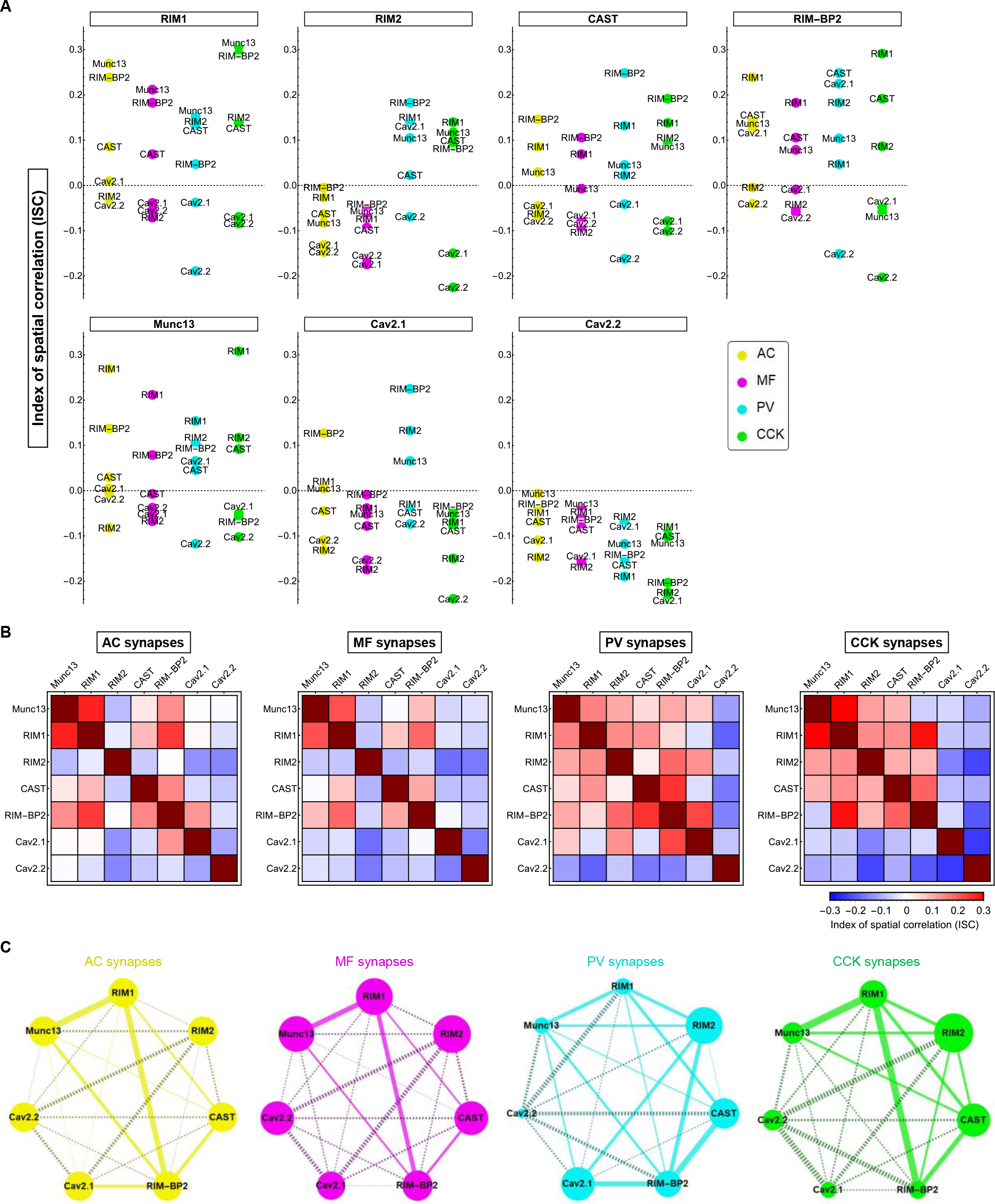
Spatial correlations between active proteins at four types of synapses. (**A**) The index of spatial correlation (ISC) between the seven active zone proteins obtained from two-color STED images (pixel size of 10 nm) are shown for the four types of synapses. Values above 0 indicate nanoscale clustering of the two proteins and, conversely, values below 0 indicate segregation of the two proteins. (**B**) Heatmap matrix of the index of spatial correlation between the seven active zone proteins for the four types of synapses. (**C**) Quantitative diagrams showing the composition and the spatial correlations of the seven active zone proteins for the four types of synapses. Dot size indicates protein density relative to AC synapses. Line thickness represents the strength of spatial correlation and solid lines depict positive clustering, while dashed lines depict negative clustering.

Surprisingly, the ISC values between RIM2 and other scaffold proteins showed positive clustering in GABAergic synapses, but segregation in glutamatergic synapses. Although the overall tendency of spatial correlation between active zone scaffold proteins was similar at the two types of glutamatergic synapses (Fig. 8B), MF synapses consistently showed lower ISC values than AC synapses.

The spatial arrangement of the synaptic vesicle release sites should relate to the efficacy of neurotransmitter release; therefore, it is particularly important to determine how Munc13 is organized into the scaffold of the active zone. The ISC value between RIM1 and Munc13 was relatively high at all types of synapses, as expected from the well-characterized interaction between Munc13 and RIM (*4, 42*), whereas the ISC value between RIM2 and Munc13 was low, especially at glutamatergic synapses (Fig. 8 A and B). Consistent with recent findings of functional coupling between RIM- BP2 and Munc13 (*43*), the ISC values between these two proteins were positive, except for at CCK synapses.

Interestingly, the ISC values between Cav2.1 and Cav2.2 were negative at all types of synapses (Fig. 8 A and B), indicating that they are well segregated at the active zone. The scaffold protein showing the highest ISC value with Cav2.1 was RIM- BP2, while the value was strongly dependent on the type of synapse: PV and AC synapses showed positive clustering of these proteins, whereas MF and CCK synapses did not. Similarly, a positive ISC value between RIM2 and Cav2.1 was only observed in PV synapses, while they were mostly segregated in AC-, MF-, and CCK synapses. None of the scaffold proteins showed positive correlation with Cav2.2. The coupling between Munc13 and Ca^2+^ channels within the active zone might determine the efficacy of neurotransmitter release (*29, 44*). We found a positive correlation between Cav2.1 and Munc13 at PV synapses, but a negative correlation at MF synapses. Although neurotransmitter release at CCK synapses is mainly triggered by Cav2.2-containing Ca^2+^ channels (*39*), Cav2.2 was segregated from Munc13 in this type of synapse as in the other synapse types.

## Discussion

We immunohistochemically quantified active zone proteins at four types of synapses, namely glutamatergic AC- and MF synapses, and perisomatic GABAergic PV and CCK synapses, in the rat hippocampal CA3 region by super-resolution fluorescence imaging techniques with simultaneous detection of pre- and postsynaptic cell-specific markers. By constructing an efficient super-resolution imaging analysis framework, we were able to systematically analyze the composition and the inter-molecular spatial relationships of seven active zone proteins, including two types of Ca^2+^ channel subunit, for thousands to tens of thousands of active zones of the four types of synapses. This revealed synapse type-specific molecular nanoconfigurations of the active zone (Fig. 8C).

### Distinct clustering of active zone proteins at multiple types of synapses

Although RIM, CAST/ELKS, and RIM-BP are redundant constituents of the active zone scaffold (*12, 13*), the composition and the inter-molecular spatial relationships of these proteins were diverse among synapse types (Figs. 7 and 8). Our analysis highlights distinctive nanotopographical motifs of scaffold proteins that associate with the release machinery of each synapse type (Fig. 8C): RIM1/RIM-BP2 forms a core scaffold that efficiently clusters Munc13 in AC and MF synapses; RIM2/RIM-BP2 forms a scaffold that clusters Cav2.1 and may couple Munc13 in PV synapses; RIM1/RIM2 forms a scaffold that redundantly clusters Munc13 in CCK synapses. We also noticed some mismatches in the clustering of active zone proteins: RIM-BP2 showed strong clustering with CAST and Cav2.1 in PV synapses, while CAST and Cav2.1 were segregated. Similarly, RIM1 showed strong clustering with RIM-BP2 and Munc13 in CCK synapses, while RIM-BP2 and Munc13 were segregated. These apparent contradictions can be explained by the clustering of active zone proteins being highly biased among active zones of the same synapse type or by the same active zone protein forming spatially distinct domains with different partners within the active zone. Whichever explanation is correct, intra-active zone compartmentalization of the active zone proteins, which seems to be more evident in GABAergic synapses, may promote functional specialization of synapses.

One finding of this study is that the density of Munc13 at the active zone differed substantially among synapse types (Figs. 7 and S5). We have previously shown that Munc13-1 forms a supramolecular nanoassembly that acts as a single release site and that the number of such assemblies per active zone corresponds to the number of release sites per active zone (*11*). The amount of Munc13 per synapse might, therefore, correlate with the number of assemblies. However, the difference in the amount of Munc13 among the different synapse types is unlikely to be caused by differences in the number of such assemblies. Rather, these differences might be attributed to differences in the number of Munc13 molecules per single release site, because the local density of Munc13 at sub-active zone hotspots varied among synapse types (Fig. 7D). Therefore, the protein stoichiometry of Munc13 in the release site assembly is not fixed but can be tuned.

The functional significance of the difference in local Munc13 density is not clear at present. Considering the diverse roles of Munc13 in synaptic vesicle priming, replenishment, and short-term plasticity (*41*), the local density might relate to the priming state of synaptic vesicles (*45*). However, the abundance of Munc13-1 alone would not determine the priming state of synaptic vesicles because its priming activity depends substantially on dimerization state; the homodimer is inactive, while the heterodimer with RIM is active (*46, 47*). Therefore, the local density of RIM would also affect overall priming activity at the release site. As revealed in this study, the two types of glutamatergic synapses have a higher ratio of Munc13 to RIM1 (Fig. S5B) and RIM2 (Fig. S5C) than perisomatic GABAergic synapses, indicating that a larger fraction of Munc13 forms inactive homodimers in the glutamatergic synapses. Furthermore, inactive spare Munc13 molecules in the supramolecular assemblies might be eligible targets of modulatory signaling molecules, including diacylglycerol (*48*), Ca^2+^/calmodulin (*49, 50*), and Ca^2+^/phospholipid (*51–53*). Thus, the high stoichiometry of Munc13 would leave plenty of room for presynaptic facilitation and potentiation at these synapses.

### Local control of Ca^2+^ channel density enables diverse release probability among synapses

In addition to PV synapses, AC and MF synapses primarily use Cav2.1 for neurotransmitter release (*54*). Among these synapse types, PV synapses show the highest release probability with a short-term depression (*39*), while MF synapses show the lowest release probability with prominent short-term facilitation (*18, 19*). Surprisingly, we found that the overall density of Cav2.1 at the active zone was not so different among these three synapse types (Fig. 7A), but we identified two Cav2.1 related factors as possible causes of the variation in release probability (PV > AC > MF synapses). First, the degree of clustering of Cav2.1 with active zone scaffolds correlated with the release probability (Fig. 8). At PV synapses, Cav2.1 positively clustered with RIM-BP2, RIM2, and Munc13, and surprisingly not with RIM1. Therefore, synaptic vesicles primed by Munc13 that interact with RIM-BP2/RIM2 should be situated in the vicinity of Cav2.1 at the active zone of these synapses. In contrast, Cav2.1 was mostly segregated from all scaffold proteins as well as from Munc13 in the active zone of MF synapses; therefore, primed synaptic vesicles should be situated away from Cav2.1. Second, the composition ratio of Cav2.1 to Munc13 correlated with release probability (Figs. 7 and S5). A recent study suggests that Munc13 interacts with RIM-BP2, which might compete with Ca^2+^ channel binding to RIM-BP2 (*43*). Thus, a high amount of Munc13 may occupy the binding sites of RIM-BP2 for Ca^2+^ channels in a supramolecular assembly of active zone proteins, thereby limiting the density of Ca^2+^ channels at the release site. Supporting this notion, the ratio of RIM-BP2 to Munc13 was also ranked in the same order. Thus, differential clustering of Munc13 might allow a wide diversity of release probability by controlling the local density of Ca^2+^channels near the release site.

Clustering of Cav2.2 largely differed from that of Cav2.1 (Fig. 8); therefore, distinct nanoscale topographical mechanisms may operate in Cav2.2 dominant synapses, such as CCK synapses. From the large differences in active zone molecular nanoconfigurations between PV and CCK synapses as well as between glutamatergic and GABAergic synapses, we infer that distinct nanoscale clustering of active zone scaffolds will diversify the release site-Ca^2+^ channel coupling among different synapse types (*55*).

We propose a model that explains synaptic diversity through synapse type-specific molecular nanoconfigurations of the active zone with the two parameters discussed above: (i) priming activity and (ii) local density of Ca^2+^ channels at the release site. MF synapses possess features matching low-release probability synapses, a wide margin for vesicle priming, and a low Ca^2+^ channel density at the release site. A sufficient margin for vesicle priming and a moderate Ca^2+^ channel density at the release site would confer the moderate release probability characteristic of AC synapses and may be requisite features for diverse synaptic plasticity, which is probably shared among cortical small glutamatergic synapses. PV and CCK synapses would show a steady priming state, while a high Ca^2+^ channel density at the release site in PV synapses would match efficient phase-locked neurotransmitter release, and a low Ca^2+^ channel density at the release site in CCK synapses, which would result from a lack of Cav2.1, should suit their distinctive asynchronous release (*39*).

Finally, we emphasize that our efficient approach opens the way to understanding causes and significance of synaptic diversity among synapses of similar types as well as among the different types of synapses. It will also be particularly useful for analyzing plastic changes of synaptic nanostructures within a large population of synapses, such as circuit-level synaptic changes induced by learning as well as by pathological alternations.

## Materials and Methods

### Animals

All experiments were performed in accordance with the policies of the Animal Ethics Committee of The University of Tokyo and The University of Yamanashi. Sprague-Dawley rats of either sex were commercially acquired from Japan SLC and housed in a group (2–4 per cage) with a 12 h light/dark cycle and food and water available *ad libitum*. Wooden or plastic toys and wood chip bedding were routinely provided.

### Antibodies

A rabbit polyclonal Munc13-1 antibody was raised against residues 1–320 of rat Munc13-1 expressed as a His-tagged recombinant protein. A rabbit polyclonal CAST antibody was raised against residues 107–138 of rat CAST expressed as a GST fusion protein. The rabbit polyclonal antibodies were affinity purified from immunized serum using the antigens. Two mouse monoclonal RIM-BP2 antibodies (clones: 8-3E IgG1 and 8-4G IgG2b) were raised against residues 621–820 of rat RIM-BP2 expressed as a His-tagged recombinant protein. A mouse monoclonal CAST antibody (clone: T-214 IgG1) was raised against a synthetic peptide of residues 107–138 of CAST. The mouse monoclonal antibodies were purified from hybridoma supernatant using protein G columns (GE Healthcare). The specificity of CAST monoclonal antibodies was confirmed by immunohistochemistry with CAST KO mice (*56*). Details of the primary antibodies used are summarized in Table S1.

All secondary antibodies were labeled with commercially available N-hydroxysuccinimide (NHS)-activated fluorescent dyes: Alexa Fluor 405, 488, 555, 594, and 647 (Thermo Fisher Scientific), Dy ight 405 and 755 (Thermo Fisher Scientific), and STAR635P (Abberior). For mouse monoclonal antibodies, highly cross-adsorbed subtype-specific secondary antibodies (IgG1, IgG2a, and IgG2b, Jackson

ImmunoResearch) were used. For rabbit and guinea pig antibodies, highly cross-adsorbed Donkey Anti-Rabbit IgG (H) and Donkey Anti-Guinea pig IgG (H) (Jackson ImmunoResearch) were used, respectively.

### Immunohistochemistry

Immunohistochemistry of rat brain sections was performed as previously described (*33*) with some modifications. Since chemical fixation often masks epitopes of active zone proteins, we precluded the use of chemical fixative (e.g., paraformaldehyde) for immunohistochemistry except in a post-fixation step. Instead, we found that flash freezing of fresh rat brain tissues with liquid nitrogen and quick dehydration of brain cryosections kept fine tissue structures intact and retained immunoreactivity of active zone proteins.

Sprague-Dawley rats of either sex (postnatal day 44–46) were deeply anesthetized with isoflurane and quickly decapitated. Brains were immediately removed from the skull and cut into slices containing the hippocampus of either hemisphere. The brain slices were then put in Tissue-Tek Cryomolds (Sakura Finetek), immersed in Tissue Freezing Medium (Leica), and instantaneously frozen on an aluminum block in liquid nitrogen. Frozen brain slices were stored in liquid nitrogen until use. For sectioning, frozen brain slices were placed in a cryostat (CM1860, Leica) at −18°C for 30 min. Then, 10 µm-thick transverse sections of the hippocampus were cut and retrieved on coverslips (Matsunami glass, 25×25 No.1, 0.13∼0.17 mm). Retrieved sections were immediately fixed by dehydration with a heat blower at 65°C for 1 min. Fixed sections were further dehydrated in ethanol at −25°C for 30 min, and then in acetone on ice for 10 min. After rehydration with PBS, sections were blocked with 0.3% BSA (Nacalai Tesque) in PBS for 1 h. Sections were then incubated with primary antibodies and then fluorescent dye-labeled secondary antibodies in PBS containing 0.3% BSA at room temperature for 3 h and 1 h, respectively. Finally, after washing off excess secondary antibodies, specimens were post-fixed with 4% paraformaldehyde (Wako) at room temperature.

### Confocal imaging

For confocal imaging experiments, specimens were immunostained with secondary antibodies bearing DyLight 405, Alexa-488, Alexa-555, and Alexa-647, respectively. Post-fixed specimens were mounted in Pro ong™ Glass Antifade Mountant (Thermo Fisher Scientific) and sandwiched with another small coverslip.

Confocal imaging experiments were performed on a Leica TCS SP8 confocal microscope equipped with a pulsed white light laser, a 405 nm diode laser, and a HyD-SMD detector. A 10× immersion objective (NA = 0.40) and a 100× oil immersion objective (NA = 1.40) were used for low magnification and high magnification imaging, respectively. All excitation laser power was set to 25% of maximum power. A HyD-SMD detector was configured to counting mode, and images were scanned at 400 Hz with 1× to 3× line accumulation. Image stacks were acquired with a step size of 250 nm. All images were acquired using Leica LAS-X software and stored as 16-bit images.

### STED imaging

For two color STED imaging, specimens were immunostained with secondary antibodies bearing Alexa-594 and STAR635P. For imaging synapse type-specific markers with confocal channels, specimens were also immunostained with secondary antibodies bearing DyLight 405 or Alexa-488. Post-fixed specimens were mounted in Pro ong™ Glass Antifade Mountant (Thermo Fisher Scientific) and sandwiched with another small coverslip.

STED imaging experiments were performed on a Leica TCS SP8 STED 3x microscope equipped with a pulsed white light laser, a continuous 592 nm STED laser (for alignment), a pulsed 775 nm STED laser, two HyD-SMD detectors, and a 100× oil immersion objective (NA = 1.40). Imaging experiments were performed at least 2 h after starting up the microscope and lasers to stabilize the system. Excitation laser wavelengths were set to 561 nm for Alexa-594 and 633 nm for STAR635P. The power of all excitation lasers was set to 25% of their maximum. The 775 nm STED depletion laser power was set to 75% to 100% of its maximum power (∼300 mW) and delay time was set to 0−300 picosecond, so that depletion coefficient reached 75−80% for Alexa594 and 85−90% for STAR635P to obtain acceptable resolution images. Alignment of the STED depletion laser was performed every 30 min. STED images of 1024 × 1024-pixel resolution were scanned at 400 Hz with 8× line accumulation. The optical zoom factor was set to 11.36, and the resulting image pixel size was 10.0 nm. Detectors were configured to counting mode with a gating from 0.5 to 6.5 ns. For comparison and confirmation of the depletion coefficient, confocal images of the same field of view (with 775 nm STED laser power 0%) were scanned prior to obtaining STED images. All images were acquired using Leica LAS-X software and stored as 16-bit images.

### STORM imaging

For two-color STORM imaging, specimens were immunostained with secondary antibodies bearing Alexa-647 405 and Alexa-647 488. Dark-red fluorescent 200 nm beads (F-8810, Thermo Fisher Scientific) were bound to the specimens and used as fiducial markers. Specimens were mounted in STORM imaging buffer, containing 50 mM HEPES (pH 8.0), 10 mM NaCl, 60% sucrose, 10% glucose, 1% β-mercaptoethanol, 0.5 mg m glucose oxidase, and 0.04 mg m catalase, and were sandwiched with another small coverslip and sealed with nail polish. High concentrations of sugar (70%) increased the refractive index of the buffer to approximately 1.47.

STORM imaging experiments were performed on a custom-built microscope with a 100× oil immersion objective (NA = 1.40, Olympus), an EMCCD camera (iXon3 860, Andor), a piezo-positioning stage (P-733.3, PI), and a custom-made objective holder that minimized mechanical sample drift during experiments (*57*). For 3- dimensional astigmatism-based calibration, a cylindrical lens (J1558RM-A, Thorlabs) was inserted into the detection path at the distance of ∼30 mm from the camera. A 640- nm laser beam (100 mW, OBIS, Coherent) passed through a Cy5 excitation filter (Semrock) was used to excite Alexa-647 molecules at an excitation intensity of ∼7.5 kW cm^2^. For detection of Alexa-647 fluorescence, a penta-band dichroic mirror (Semrock) and a bandpass emission filter (710 M80, Omega) were used. Raw image data of 128 × 128-pixel resolution were obtained as 16-bit tiff image stacks. The raw image pixel size was 200 nm.

Laser beams of 405 nm and 488 nm (OBIS, Coherent) were used to activate Alexa-647 molecules via Alexa-405 and Alexa-488, respectively. A series of 10,000 imaging frames (10 ms per frame) were alternately measured for 488 and 405 channels for a total of 120,000 imaging frames for image reconstructions. The excitation intensities of the 488 nm and 405 nm lasers were gradually increased from 0.25 to 8 W cm^2^ and from 1 to 32 W cm^2^, respectively, so that fluorescent signals from activated molecules were not spatially overlapped.

Localizations of fluorophores were determined using custom-built software in ImageJ. A 2D Gaussian model was used with the linear least-square fitting method. Z-axis localizations were determined using 3D astigmatism-based calibration. Sample drift during recording was corrected using fluorescent fiducial markers. STORM images were reconstructed by pixelating Alexa-647 localization datasets with a 10 nm binning as 32-bit tiff images and processed using a 3D Gaussian spatial filter with a radius of 4 pixels (40 nm).

### STED imaging analysis

All data analysis was performed using Mathematica software version 12.0.0 (Wolfram). For representation and quantification of two-color STED imaging results (Alexa-594 and STAR635P channels), raw STED images (pixel size of 10 nm) were deconvolved using a gaussian kernel with a radius of 4 pixels (40 nm). The ambient background (non-synaptic) fluorescent signals in STED images were locally calculated from the non-synaptic region surrounding synaptic sites, which were estimated using an unsharp masking method (see below), and then subtracted from deconvolved STED images for each channel.

To obtain image masks for the active zones in the systematic nanoscopic image processing pipeline (Fig. 6), merged two-color STED images of active zone proteins were processed by unsharp masking. In this process, STED images filtered by gaussian blur with a radius of 40 pixels (400 nm) were subtracted from STED images filtered by gaussian blur with a radius of 4 pixels (40 nm). The resultant subtracted images were then binarized, and components larger than 100 pixels (0.01 µm^2^) were included in the following analysis as reasonable estimates for the area of active zones. Selected active zones were classified into glutamatergic- and GABAergic synapses by co-localization with a binarized mask of synapse type-specific markers (PSD95 and gephyrin, respectively) imaged using a confocal channel. An additional binarized mask of a presynaptic marker (Syt2 or VGLUT3) imaged using another confocal channel was used to further categorize GABAergic synapses into PV- and CCK synapses. The mean pixel intensities of respective STED channels and pixel-by-pixel Pearson’s correlation coefficients between two-color STED channels were calculated using the obtained masks as estimates of protein density (Figs. 7A and S5) and the spatial correlation between the pair of proteins (Fig. S6), respectively.

Image masks for sub-active zone hotspots of active zone proteins were separately generated from respective STED channels by combining image deconvolution and unsharp masking. In this process, STED images filtered by gaussian blur with a radius of 4 pixels (40 nm) were subtracted from STED images deconvolved using a gaussian kernel with a radius of 4 pixels (40 nm) to detect the edges of small hotspots of the active zone proteins. The resultant images were then binarized, and components larger than 10 pixels (0.001 µm^2^) were included in the analysis. The obtained masks for sub-active zone hotspots were used to quantify the number (Fig. 7B) and size of hotspots (Fig. 7C), the local density of the protein (Fig. 7D), and the degree of co-localization (Fig. S7). To compare the degree of co- localization with Pearson’s correlation coefficients (which range from -1 to 1), the slope values in Fig. S7 were transformed using a hyperbolic tangent function as the index of the degree of colocalization (Fig. S8).

### Data and materials availability

All data, custom-written code files, and materials that support the findings of this study are available from the corresponding author upon reasonable request.

## Ac nowledgments

We thank Dr. T. Onishi and Dr. M. Kono for technical assistance and helpful discussions in developing the immunohistochemical method, Dr. A. Hagiwara and Dr. Y. Hida for technical support in producing the CAST antibody. We appreciate the support of The IRCN Imaging Core, The University of Tokyo Institutes for Advanced Studies. This work was supported by Grants-in-Aid for Scientific Research (KAKENHI) from the Ministry of Education, Culture, Sports, Science, and Technology of Japan (MEXT) (19K16251, 20KK0171, and 21K15183 to H.S., and 19K22247 and 20H03427 to K.H.), JST, PRESTO, Japan (Grant Number JPMJPR21E7 to H.S.) and JST, CREST, Japan (Grant Number JPMJCR1751 to T.O.).

## Author contributions

H.S. conceived the project, performed the experiments, analyzed the data, and wrote the manuscript. N.K. and S.N. produced monoclonal antibodies for RIM-BP2. S.H. and

T.O. produced a monoclonal antibody for CAST. K.H. supervised the study and discussed the results. H.S. and K.H finalized the manuscript.

---

**Figure S1.**
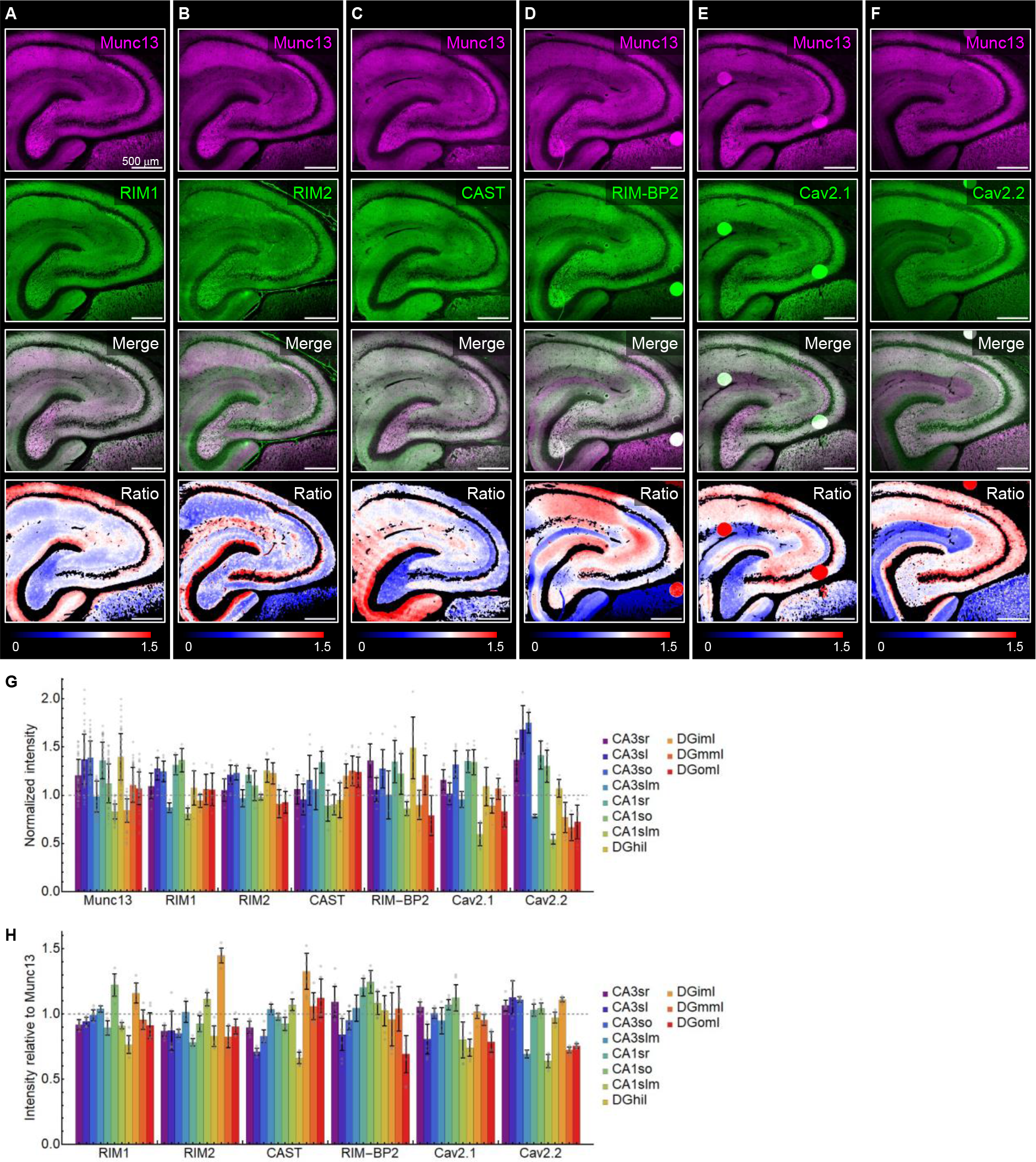
Immunohistochemical analysis of active zone proteins in the hippocampus. (**A**–**F**) Immunofluorescence images of Munc13 (magenta) and active zone proteins (green) in the hippocampus. The bottom panels show the ratio image between active zone proteins and Munc13. Scale bars, 500 µm. ( ) uantification of the fluorescence intensity of the seven active zone proteins in various sub-regions of the hippocampus. sr: stratum radiatum; sl: stratum lucidum; so: stratum oriens; slm: stratum lacunosum- moleculare; hil: hilus; iml: inner molecular layer; mml: middle molecular layer; oml: outer molecular layer. The data were normalized against the overall mean values for each protein. Error bars show standard deviations. (**H**) uantification of the relative fluorescence intensity of active zone proteins to Munc13 in various sub-regions of the hippocampus.

**Figure S2.**
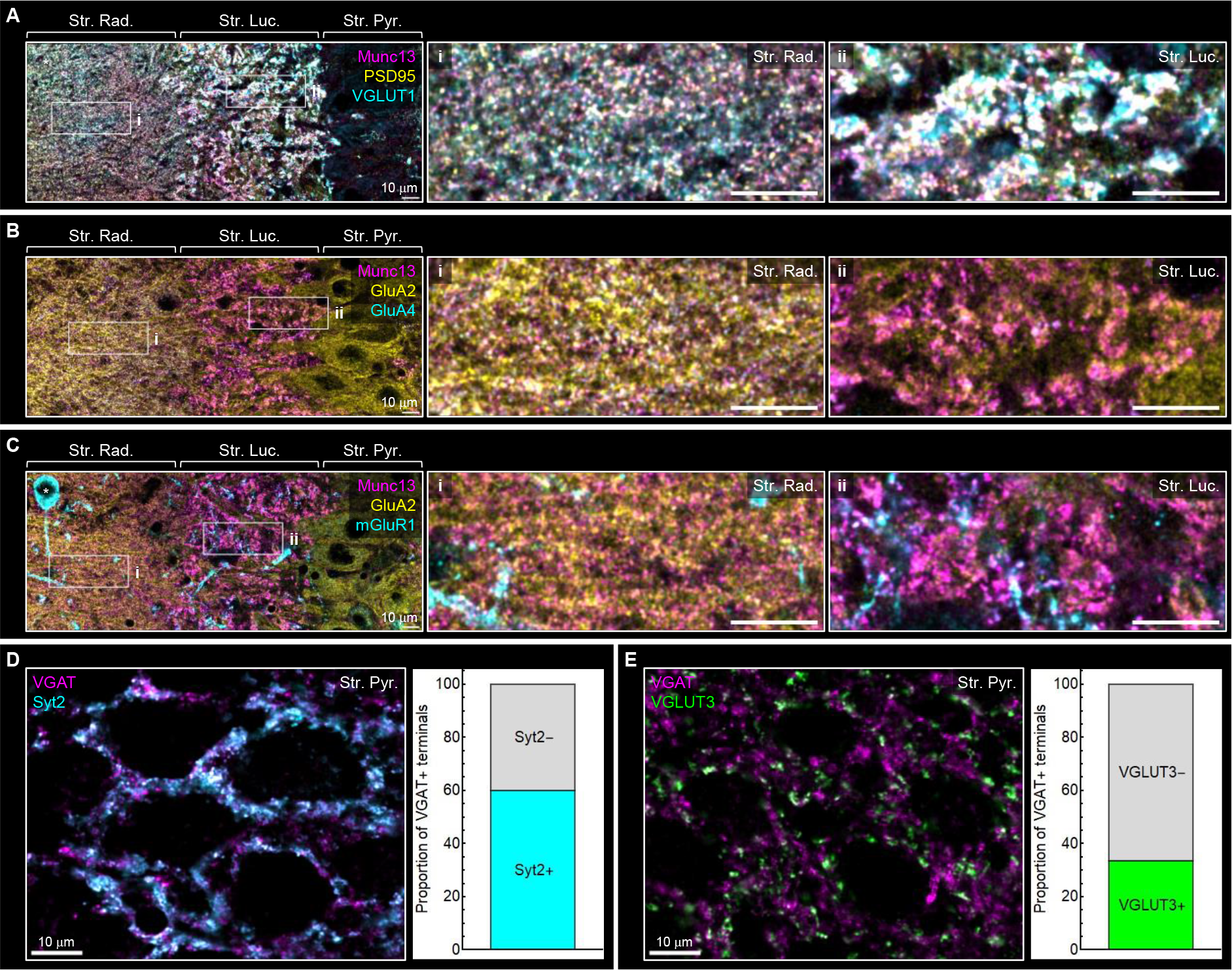
Characterization of synapses by immunofluorescence of pre- and postsynaptic markers in the hippocampal CA3 region. (**A**–**C**) Three-color confocal images of Munc13 (magenta) and two other synaptic proteins (yellow and cyan): PSD95 and G UT1 (A); GluA2 and GluA4 (B); GluA2 and mGluR1α (C). The right two panels show enlarged views of the stratum radiatum (i) and the stratum lucidum (ii) indicated by the white boxed regions in the left panel. Scale bars, 10 µm. (**D** and **E**) Two-color confocal images of VGAT and Syt2 (D) or VGLUT3 (E). The bar graphs depict quantification of the ratio of GABAergic synapses positive for Syt2 (D) and VGLUT3 (E) in the stratum pyramidale of the hippocampal CA3 region. Scale bars, 10µm.

**Figure S3.**
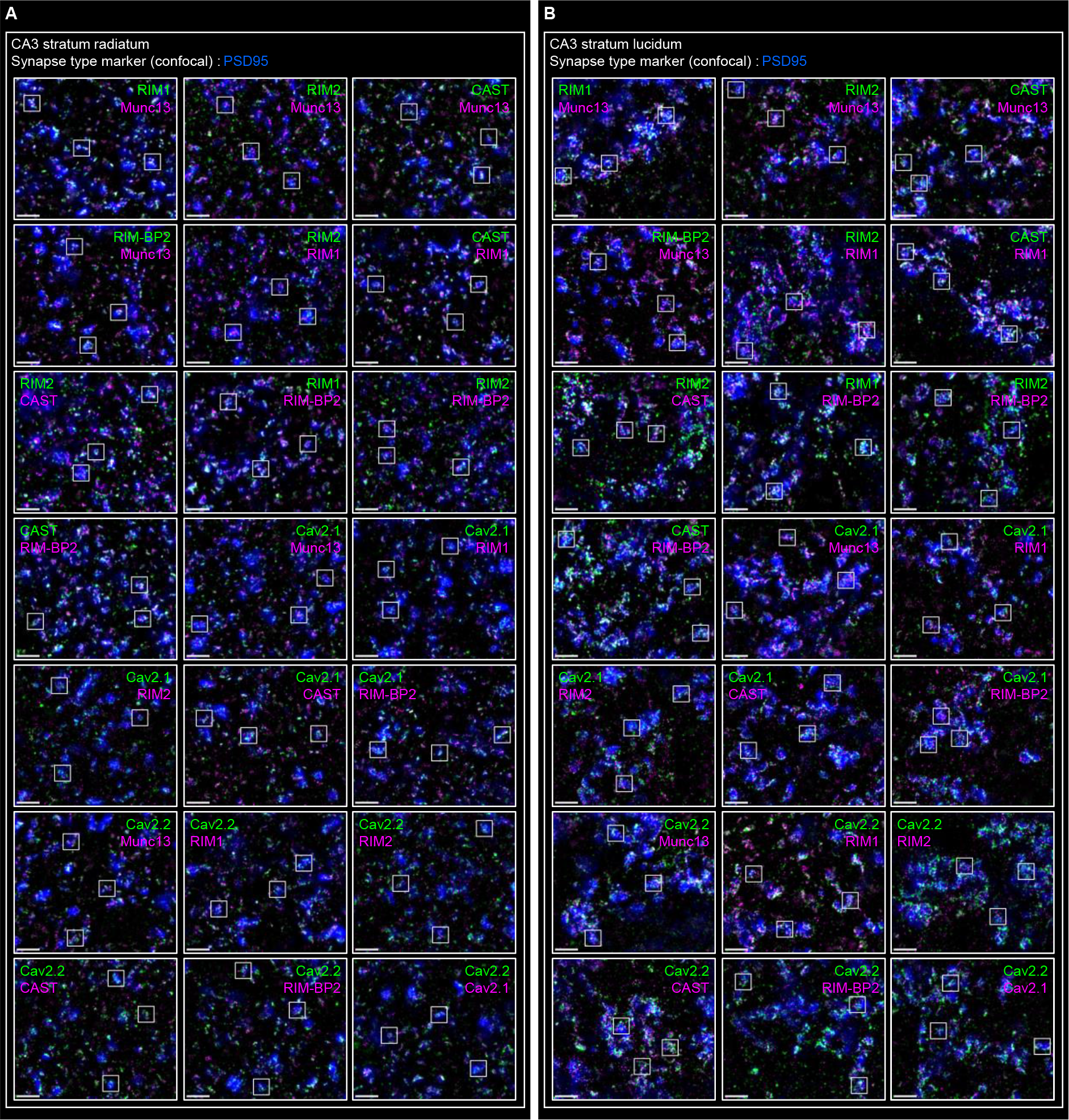
Systematic nanoscopic imaging of active zone proteins at glutamatergic synapses in the hippocampal CA3 region (related to Fig. 4). (A **and B**) Two-color STED images of active zone proteins (green and magenta) overlaid with confocal images of a synaptic marker for glutamatergic synapses (PSD95; blue) in the stratum radiatum (A) and the stratum lucidum (B) of the hippocampal CA3 region. Scale bars, 500 nm. Enlarged views of the white boxed regions are shown in Fig. 4.

**Figure S4.**
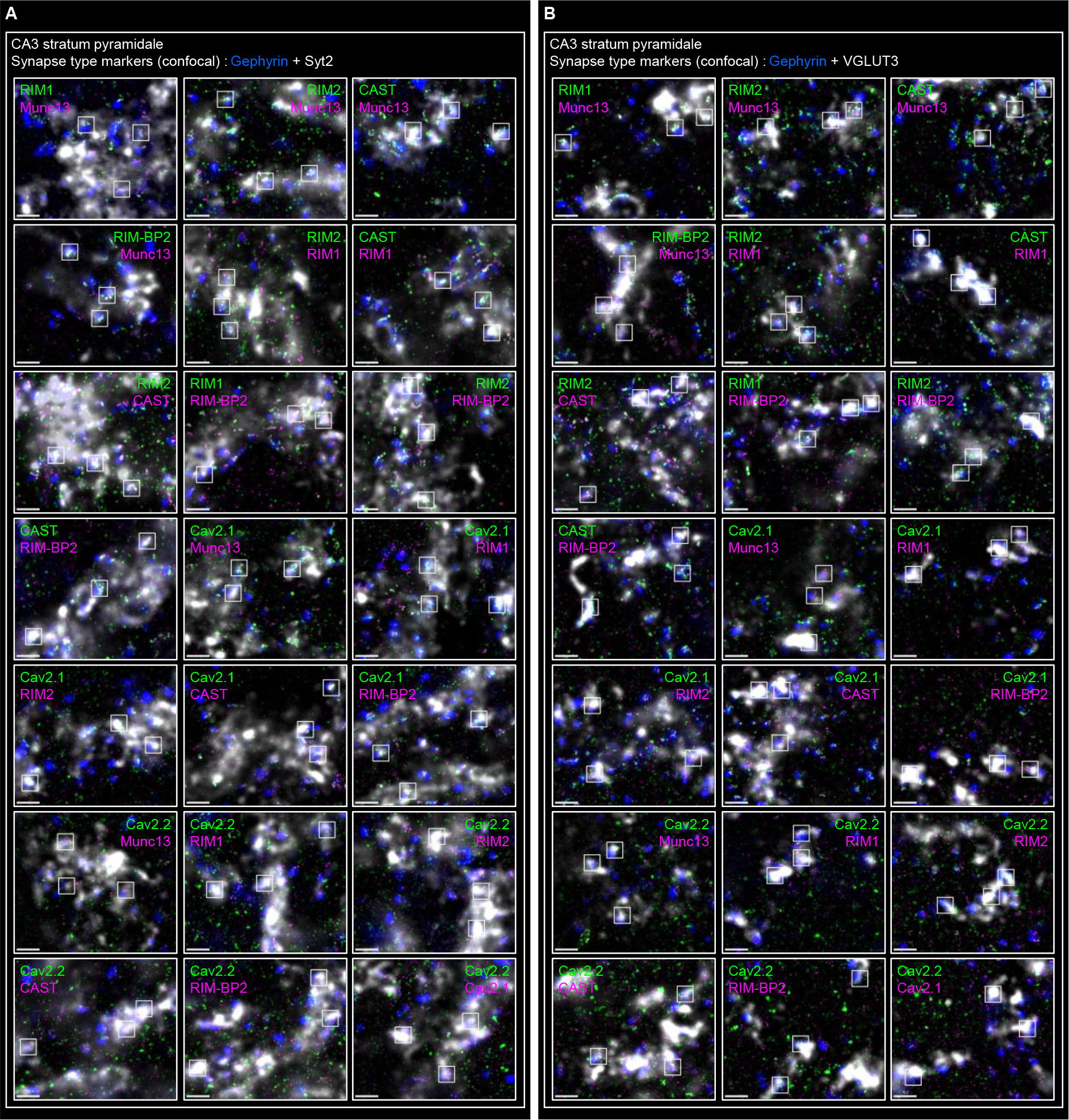
Systematic nanoscopic imaging of active zone proteins at perisomatic ABAergic synapses in the hippocampal CA3 region (related to Fig. 5). (**A** and B) Two-color STED images of active zone proteins (green and magenta) overlaid with confocal images of a synaptic marker for GABAergic synapses (gephyrin; blue) and a synapse-type marker (white) in the stratum pyramidale of the hippocampal CA3 region. Syt2 (A) and G UT3 (B) are synapse-type specific markers that identify P and CCK synapses, respectively. Scale bars, 500 nm. Enlarged views of the white boxed regions are shown in Fig. 5.

**Figure S5.**
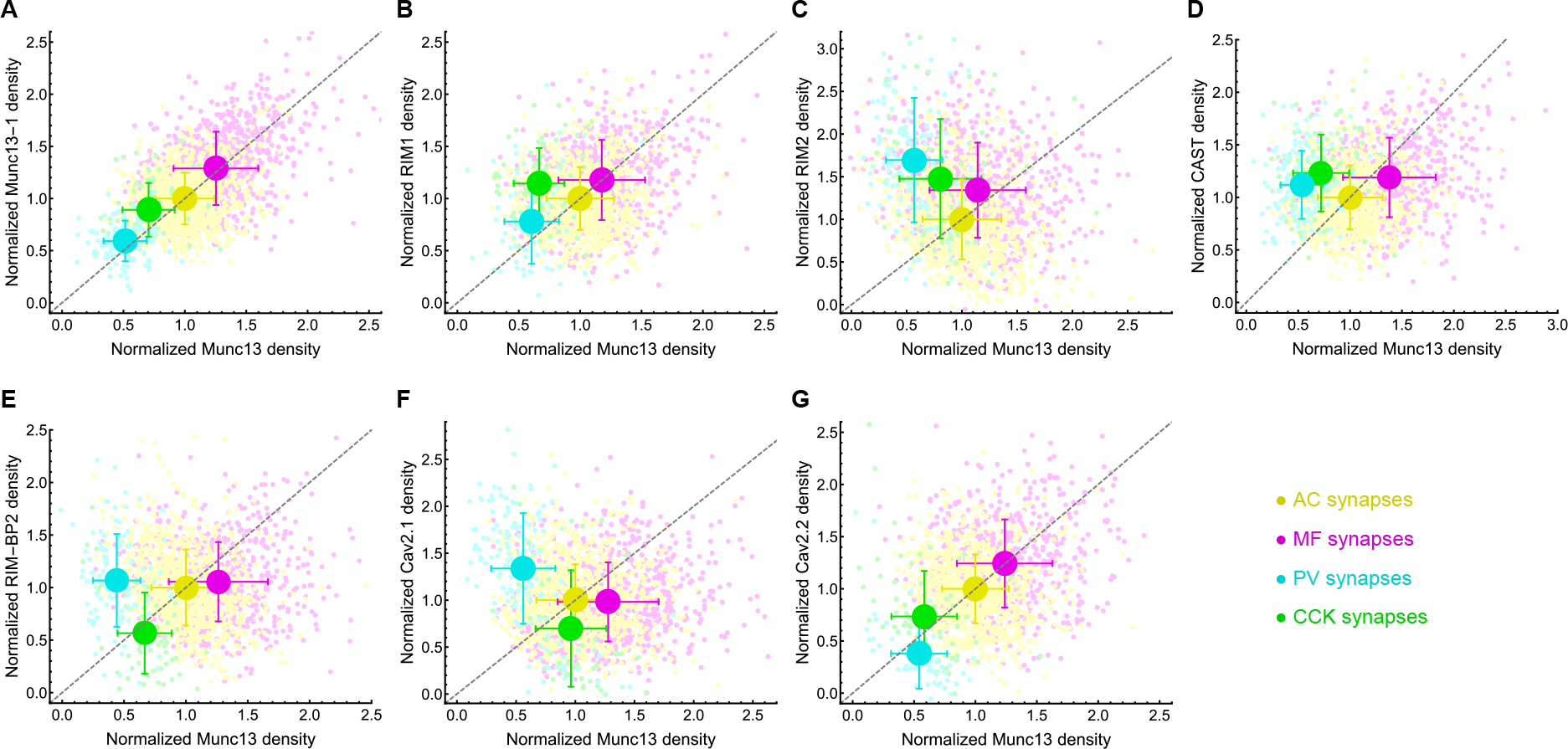
The density of active zone proteins relative to Munc13. (**A**–**G**) The density of Munc13-1 (A), RIM1 (B), RIM2 (C), CAST (D), RIM-BP2 (E), Cav2.1 (F), and Cav2.2 (G) is plotted against that of Munc13 at individual active zones for the four types of synapses. Large dots represent the mean values for each synapse type. Error bars show standard deviations. The value for each protein is normalized against the mean value of AC synapses .

**Figure S6.**
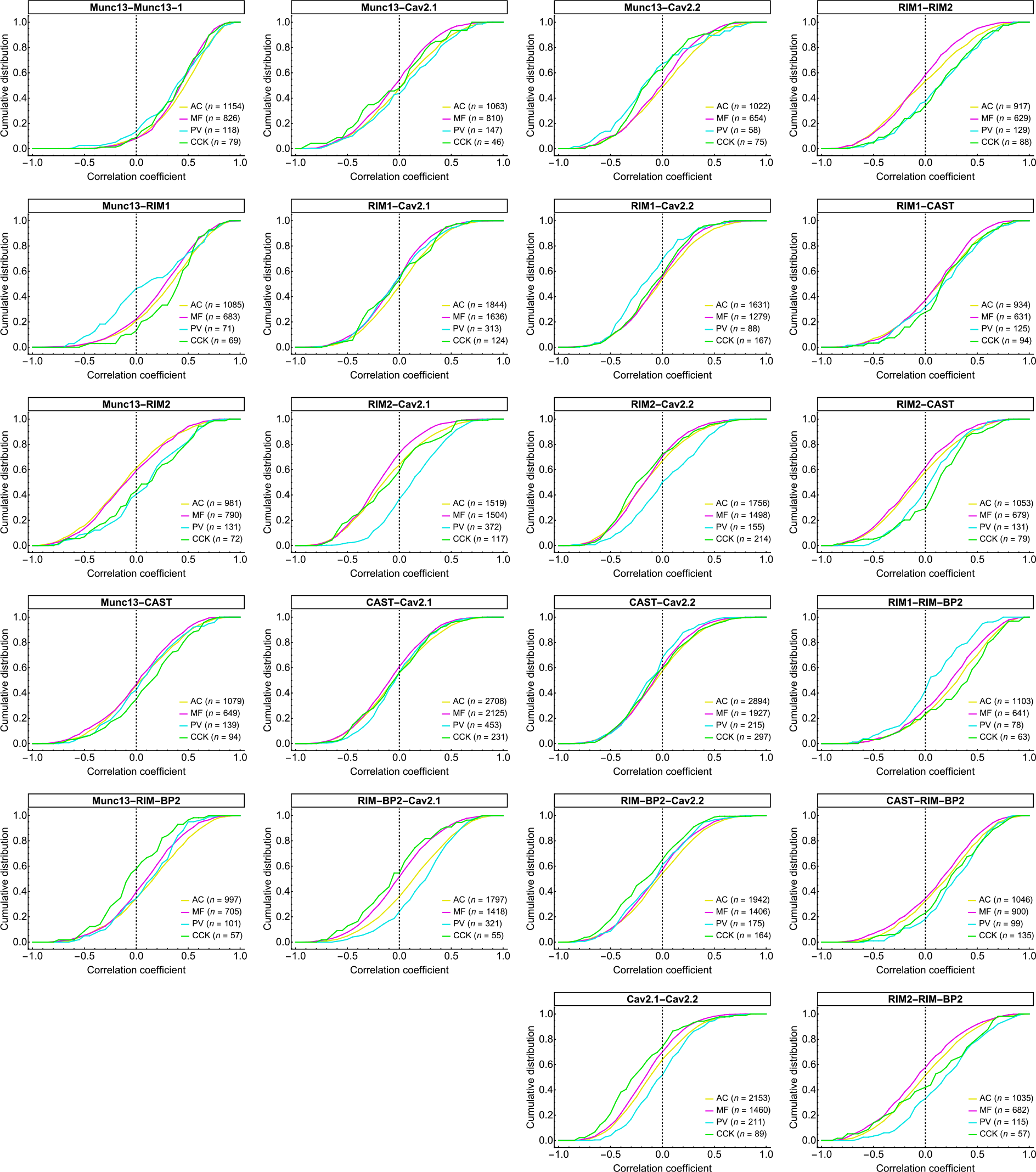
Nanoscale spatial correlation of active zone proteins. Pixel-by-pixel correlations (Pearson’s correlation coefficients) between two-color STED images (pixel size of 10 nm) for the seven active zone proteins are quantified at individual active zones for the four types of synapses.

**Figure S7.**
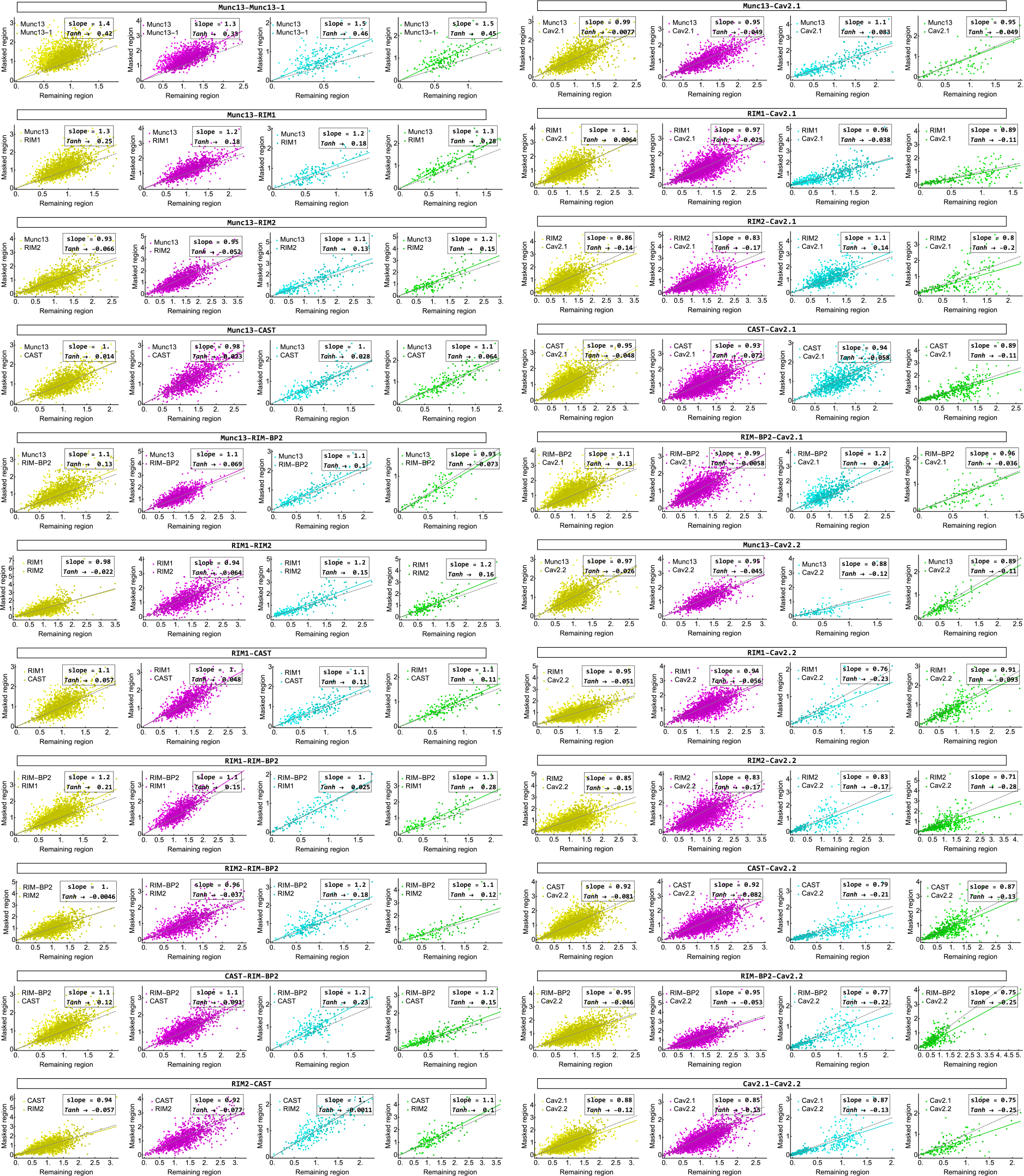
Nanoscale co-localization analysis of active zone proteins. The mean pixel intensities of one of the two STED image channels within and outside of the hotspot mask defined by the other channel (see Fig. 6) are calculated for each active zone and displayed as scatter plots for the four types of synapses (yellow: AC synapses; magenta: MF synapses; cyan: PV synapses; green: CCK synapses). The values are normalized against the overall mean pixel intensity at active zones of AC synapses. The straight lines are the least square fits to the data. A slope greater than 1 indicates colocalization of the two proteins within the active zone, while a slope less than 1 indicates segregation.

**Figure S8.**
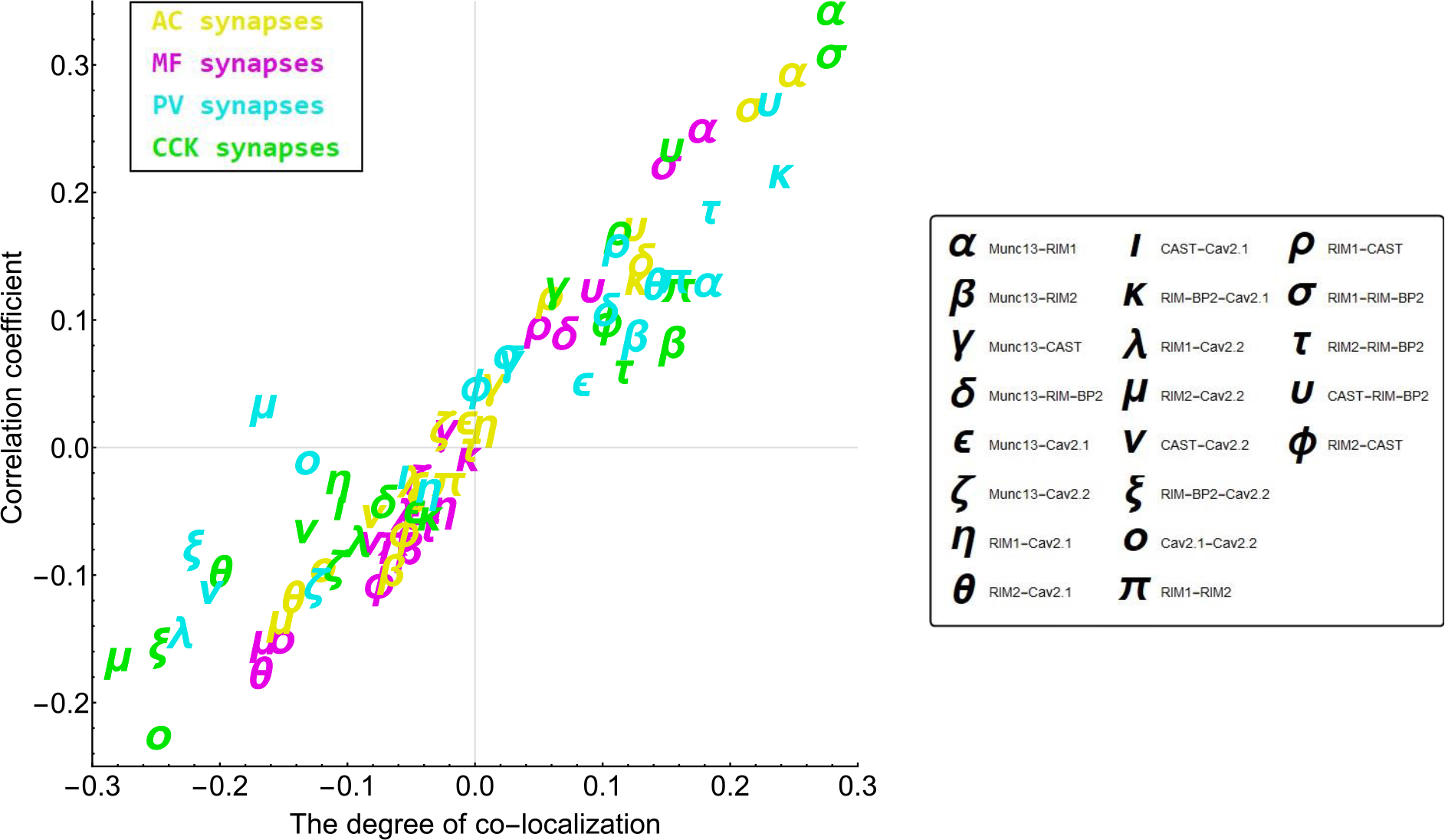
Relationship between two types of independent estimate of nanoscale spatial correlation of active zone proteins. The degree of colocalization obtained from the analysis of two-color STED images in Fig. S7 is plotted against the Pearson’s correlation coefficients obtained from the analysis in Fig. S6 Greek letters indicate protein pair combinations for the seven active zone proteins. These two types of independent estimates of nanoscale spatial correlation were well matched.

**Table S1.**
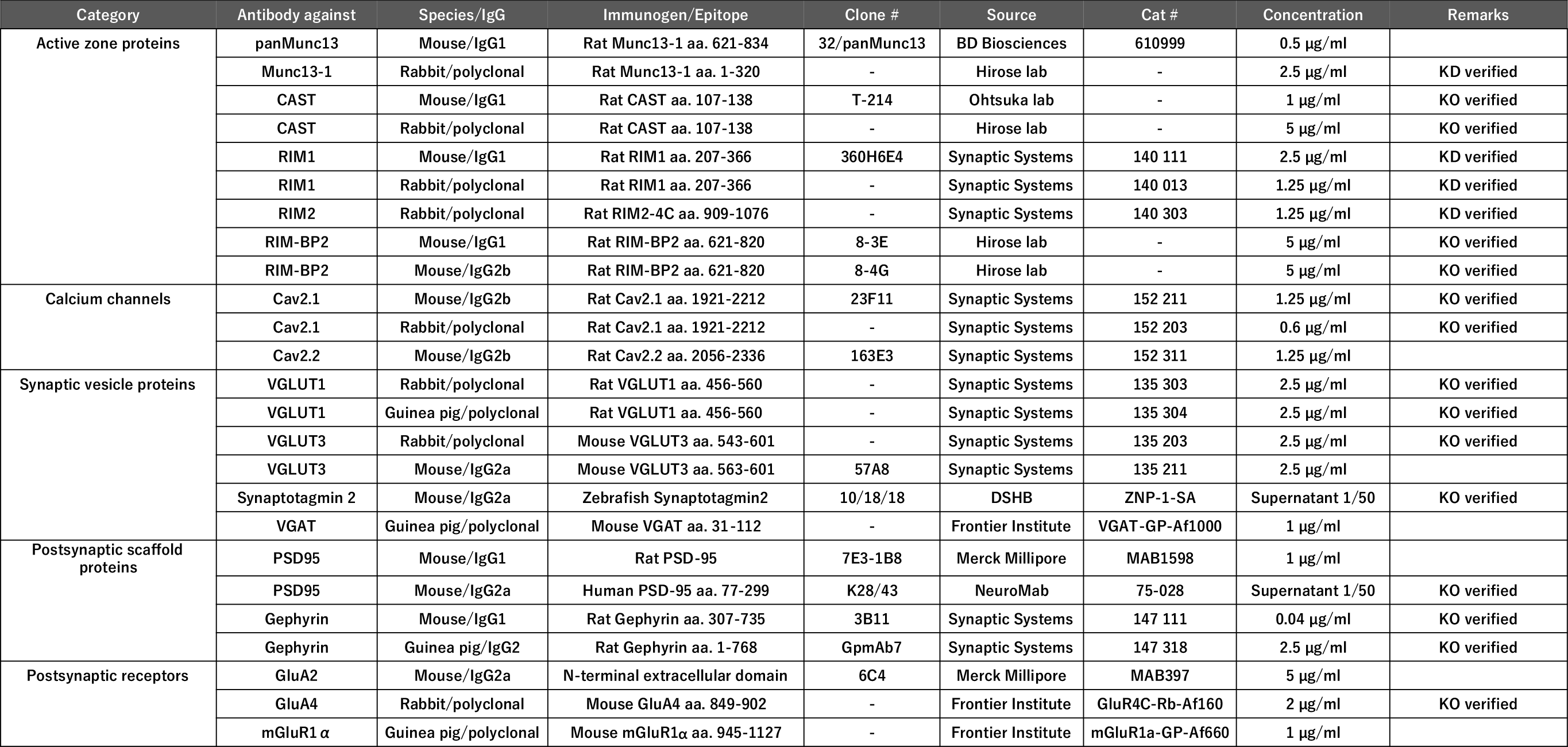
Summary table of primary antibodies used in this study

